# Simulated 5-HT2A receptor activation accounts for the high complexity of brain activity during psychedelic states

**DOI:** 10.1101/2025.10.20.683366

**Authors:** Hugo M. Martin, Rodrigo Cofre, Alain Destexhe

## Abstract

Serotonergic psychedelics, such as LSD, psilocybin, and DMT, have strong effects on human brain activity, yet their mechanisms of action are only partially understood. Here, we present a biophysically-based mean-field model that integrates cellular and network-level details to simulate the effects of these compounds at different spatial scales. By incorporating the brain-wide distribution of 5-HT2A receptors, our model mechanistically links receptor activation through reducing leak membrane potassium conductances, based on electrophysiological data. Our simulations reveal that this microscopic perturbation leads to the emergence of a brain state characterised by asynchronous irregular dynamics with increased firing rates and alterations in spectral power, in particular reduction of alpha-frequency bands, consistent with empirical findings. This change in dynamics is accompanied by an increase in spontaneous complexity, as quantified by the Lempel-Ziv complexity index, as observed experimentally. Furthermore, our model accurately replicates experimental findings regarding the Perturbational Complexity Index (PCI), demonstrating that PCI does not increase significantly by psychedelic drug administration. This crucial dissociation, where spontaneous complexity and spectral power are affected while perturbational complexity is preserved, highlights the distinct neurophysiological substrates underlying different metrics in psychedelic states. Our multiscale model provides a robust, mechanistic framework for understanding how psychedelics modulate global brain activity.

## 1 Introduction

Over the past few years, research on psychedelics has gained significant momentum, driven predominantly by their therapeutic potential in treating mood disorders and addictions [1, 2, 3]. This growing interest emphasizes the need to mechanistically understand the effects of psychedelic compounds on the brain and the need to integrate in biophysical models consistent findings from the experimental literature.

Recent advances in neuroimaging have provided novel insights into brain organization across multiple scales, enabling the development of computational models that integrate biophysical mechanisms with large-scale neural dynamics [4, 5, 6, 7, 8]. These whole-brain models provide a rich tool for identifying robust biomarkers of different states of consciousness, offering a principled framework to investigate how changes at the molecular and cellular scales translate into macroscopic patterns of brain activity [9]. In this context, modeling the action of serotonergic psychedelics at the whole-brain level provides a unique opportunity to explore the dynamical principles underlying consciousness alterations, allowing the integration of different neuroimaging data such as functional and diffusion MRI and PET scans [10, 11, 12, 13, 14, 15].

The action of serotonergic psychedelics in the brain originates at the molecular and cellular levels, where receptor binding and intracellular signaling cascades drive alterations in neuronal excitability and network activity [16, 17, 18, 19]. The integration of molecular and cellular details into whole-brain models remains computationally prohibitive, emphasizing the need for theoretically grounded frameworks that link molecular scale interactions to whole-brain neural dynamics.

In this work, we use a computational framework designed to bridge the gap between molecular and cellular processes and whole-brain dynamics, addressing the challenge of biologically plausible large-scale brain simulations. At the heart of our approach is a biophysically principled meanfield model that incorporates cellular and molecular mechanisms while reducing the system’s dimensionality to enable computationally efficient large-scale simulations. By capturing emergent network properties across multiple spatial scales, our model enables realistic whole-brain simulations of serotonergic psychedelic action. Our hierarchical modeling approach is based on a previously published framework [9], that consists of four interdependent levels: (i)single-cell models that describe molecular interactions and intrinsic neuronal dynamics (microscale);(ii)networks of spiking neurons that characterise local circuit activity (mesoscale); (iii) a mean-field model that approximates collective network behavior while maintaining biophysical accuracy (mesoscale, low-dimensional); and (iv) a whole-brain model leveraging mean-field dynamics to simulate large-scale neural activity constrained by empirical anatomical connectomes (macroscale). To implement and validate our simulations, we employ The Virtual Brain (TVB) platform [20].

Specifically, here we present a bottom-up model of serotonergic psychedelic action that explicitly incorporates the effects of reduced leak potassium (K+) conductance. The proposed mean-field model integrates membrane conductances and synaptic receptor dynamics to simulate serotonergic modulation mediated by 5-HT_2A_ receptors, which are considered the principal targets through which serotonergic psychedelics elicit hallucinogenic effects in humans [21, 22]. Activation of these receptors has been shown to modulate leak K+ conductances, consistent with electrophysiological evidence [23, 24, 25, 26, 27, 28].

At the whole-brain scale, psychedelic compounds consistently increase spontaneous signal diversity in the human brain, a neural marker associated with altered conscious content. Studies using electroencephalography (EEG) and magnetoencephalography (MEG) have shown increased neuronal complexity in a variety of psychedelic states. These effects are robustly captured by metrics such as Lempel-Ziv complexity (LZc) and are frequently accompanied by reductions in delta-, theta- and alpha-band oscillatory power [29, 30, 31, 32, 33, 34, 35].

Importantly, such increases in spontaneous complexity do not necessarily translate to changes in evoked cortical dynamics. Studies employing transcranial magnetic stimulation coupled with EEG (TMS–EEG) report that while psychedelics such as psilocybin and ketamine enhance spontaneous signal diversity, the Perturbational Complexity Index (PCI)—an index of the brain’s causal capacity to integrate and differentiate information—does not increase signifcantly [32, 31]. This dissociation suggests that spontaneous complexity may reflect the richness of conscious content, whereas evoked complexity may be more tightly linked to the underlying capacity for consciousness itself.

Incorporating receptor density distributions across the brain, our model predicts that serotonergic psychedelics induce an activated brain state characterised by asynchronous-irregular dynamics and increased complexity relative to the waking resting state, linking synaptic receptor activity to whole-brain network dynamics.

Consistent with empirical findings, we observe our model exhibits heightened brain complexity, as measured by LZc. Furthermore, our simulations of evoked activity show that, in agreement with experimental data, PCI increases, but not significantly under psychedelics. We also observe a robust decrease in spectral power within the *δ, θ*, and *α* bands under reduced K+ conductance, consistent with the literature on serotonergic psychedelics.

## 2 Results

This section introduces the framework that connects single-neuron excitability to large-scale activity. We then show how the decreased potassium leak conductance increases neural excitability and, in turn, drives measurable alterations in brain state, reflected in spontaneous complexity (LZc), evoked complexity (PCI), and spectral power.

### 2.1 Multiscale Modeling Framework from Single Neurons to Whole-Brain Dynamics

To investigate how cellular-level mechanisms shape whole-brain dynamics under serotonergic modulation, we implemented a multiscale modeling framework spanning single neurons, local circuits, and the macroscale connectome (Figure 2).

**Figure 1:**
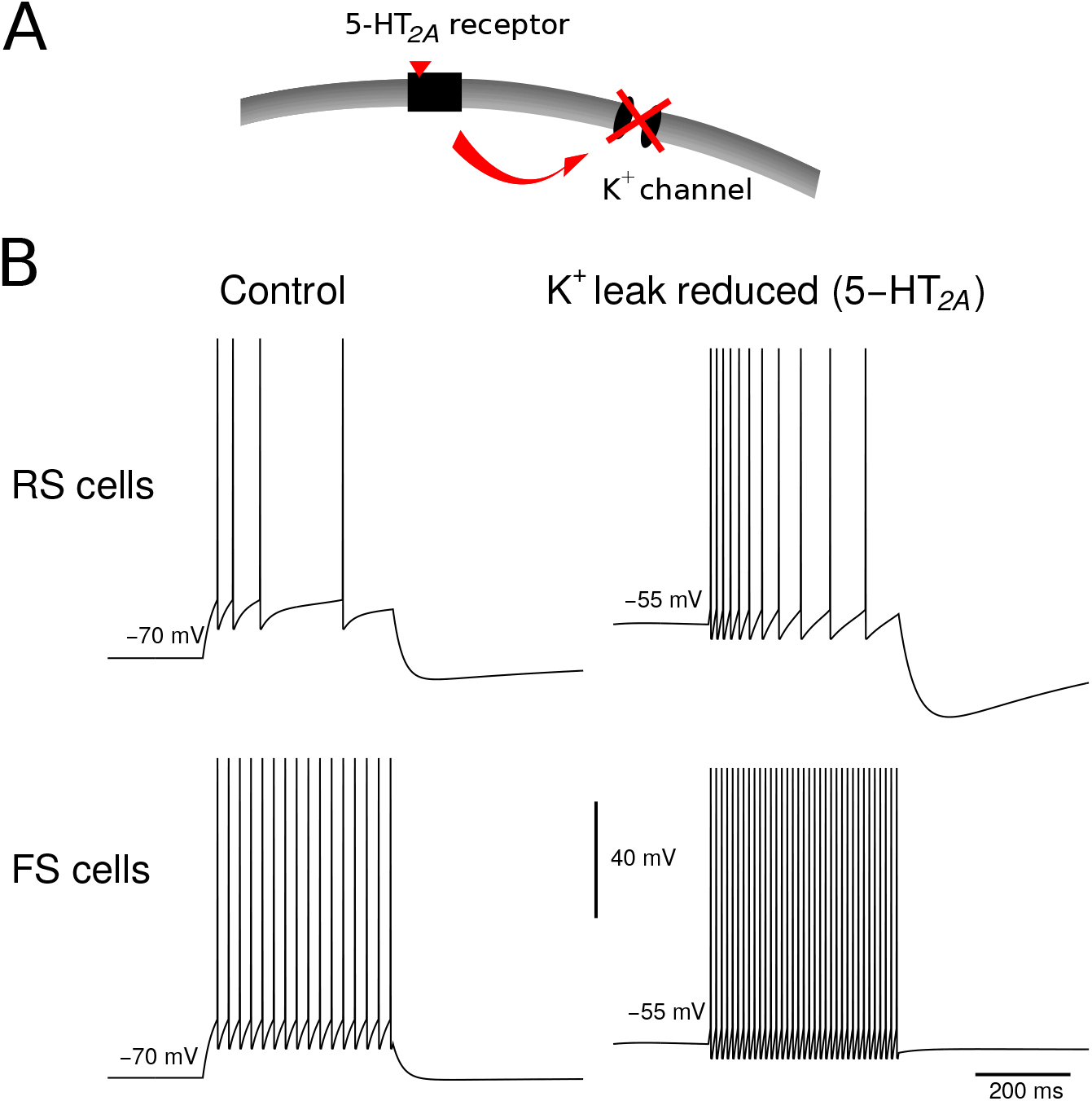
5-HT_2A_ receptor activation depolarizes cortical neurons by reducing K+ leak conductance. **(A)** Schematic illustrating the biophysical mechanism where 5-HT_2A_ receptor activation closes potassium (K+) leak channels, reducing outward current and leading to depolarization. **(B)** Electrophysiological traces from regular-spiking (RS, top) and fast-spiking (FS, bottom) cortical neurons in control conditions (left) and following simulated 5-HT_2A_ receptor activation via reduced K+ leak conductance (right). For both cell types, reducing K+ leak conductance from 8.21 to 5.37 nS depolarizes the resting membrane potential from −70 to −55 mV and increases firing rate in response to the same current injection. This models the excitatory effect of serotonergic modulation mediated by 5-HT_2A_ receptors.

**Figure 2:**
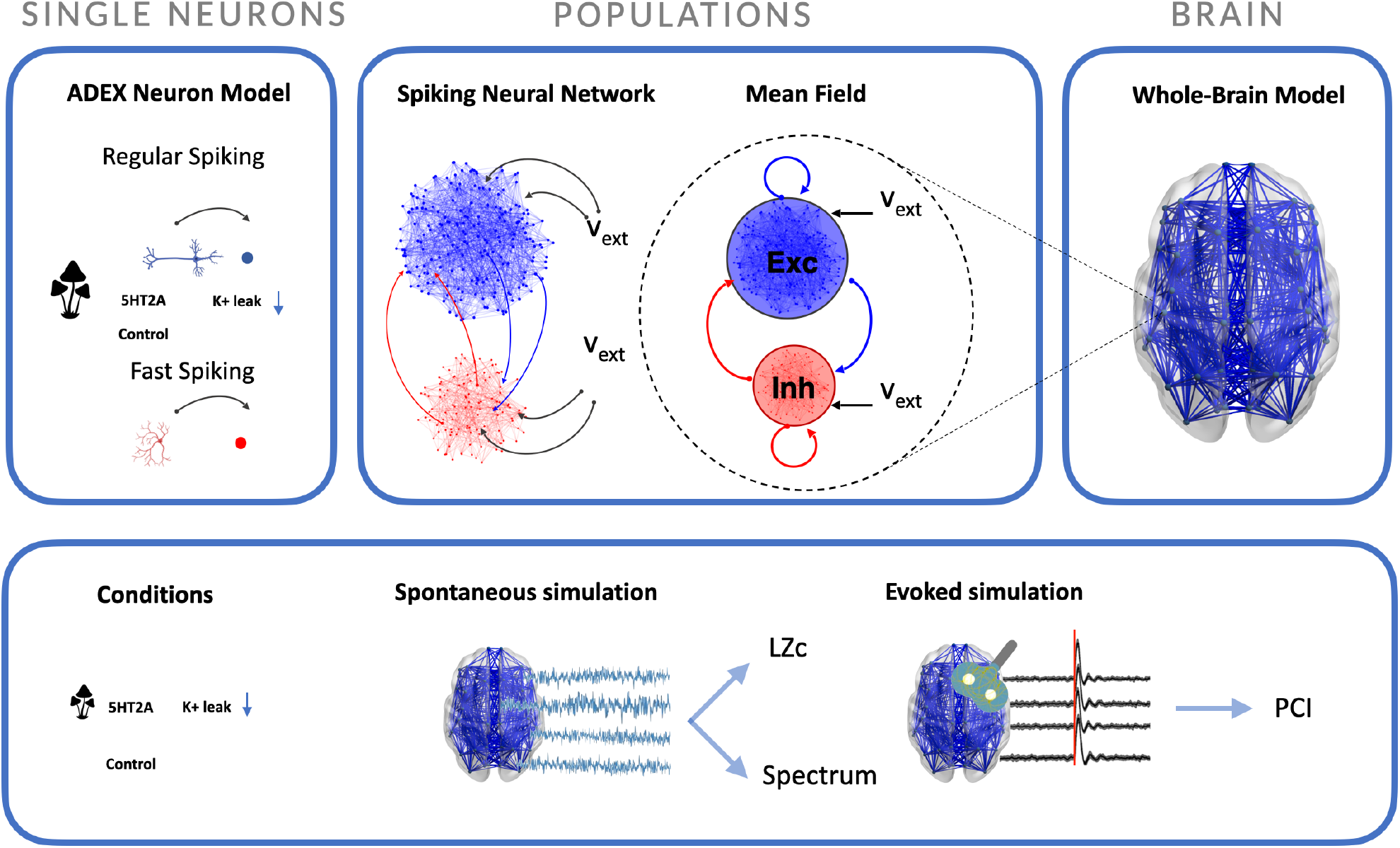
Multiscale Modeling Framework for Brain Dynamics. Our modeling framework integrates neural activity across three scales: single neurons, local populations, and the whole-brain. **Top row:** At the single neuron scale, we used the Adaptive Exponential (AdEx) integrate-and-fire model, because its capacity to reproduce diverse firing patterns such as regular spiking and fast spiking, influenced by factors like 5-HT_2A_ receptor activation and K+ leak currents. The population scale illustrates two complementary approaches: a detailed spiking neural network (left), depicting excitatory (blue) and inhibitory (red) populations with external inputs (*V*_*ext*_), and a mean-field model (right), showing the mean dynamics of excitatory (Exc) and inhibitory (Inh) populations and their interactions. At the whole-brain scale, the model is built by connecting multiple mean-field units based on anatomical connectivity. **Bottom row:** The framework’s utility in simulating different conditions (e.g., control and 5-HT_2A_ activation) under both spontaneous and evoked simulation paradigms. Spontaneous activity can be analysed for metrics such as Lempel-Ziv complexity (LZc) and spectral properties. Evoked simulations are used to analyse responses to external stimuli, leading to the quantification of the Perturbational Complexity Index (PCI). This hierarchical approach enables the investigation of how cellular-level mechanisms give rise to emergent macroscopic brain states.

At the microscale, we employed the Adaptive Exponential integrate-and-fire (AdEx) model, balancing biological plausibility with computational tractability in large-scale simulations [36, 37]. The AdEx model captures a wide repertoire of cortical firing behaviors, including regular spiking and fast spiking. The model is governed by two coupled differential equations: one describing membrane potential dynamics (Eq.7) and another describing the adaptation current (Eq.8), which regulates neuronal excitability as a function of spiking history. Different parameterizations allow the representation of excitatory pyramidal neurons and inhibitory interneurons, fitted from in vitro electrophysiological recordings [38]. This versatility permits adaptation to multiple brain structures, including the thalamus, basal ganglia, and cerebellum [39, 40, 41].

At the mesoscale, we constructed networks of spiking neurons to represent local microcircuits or anatomically defined regions. In a canonical cortical column model, 10,000 neurons with an 80:20 ratio of pyramidal cells to interneurons which are randomly connected with probability *p* = 0.05, consistent with empirical estimates [42]. Synaptic transmission was modeled as conductance-based via AMPA/NMDA (excitatory) and GABA_A_ (inhibitory) receptors.

To efficiently bridge from high-dimensional spiking networks to population dynamics, we employed a mean-field approach derived from a second-order Master Equation [43]. This approach preserves key biophysical features—such as mean activity, conductance fluctuations, and adaptation—while enabling rapid simulation of large-scale systems. The core of this method is the neuronal transfer function, which relates firing rate output to presynaptic input. For conductance-based models without closed-form solutions, we use a semi-analytical procedure: numerically simulating neuron responses across input conditions and fitting them with analytical templates, validated for AdEx neurons [44, 38] and extended to more complex neuron types s such as the quadratic integrate-and-fire or Hodgkin–Huxley formulations [45, 46, 41].

At the macroscale, the brain was modeled as a network of mean-field units coupled according to empirically derived structural connectivity. The chosen parcellation defines the model’s spatial resolution. We used the 68-region Desikan–Killiany atlas [47] for the results in the main article, but we replicate all our results using the cortically-reduced Lausanne 463 parcellation. Inter-regional coupling strengths and conduction delays were derived from diffusion MRI tractography, with delays computed from anatomical distances. Each regional unit receives both stochastic background input, modeled as Ornstein–Uhlenbeck process, and weighted, time-delayed inputs from other regions.

Figure 2 illustrates this hierarchical architecture. The top row shows the three scales: single-neuron AdEx models reproducing diverse firing patterns and modulations, including those induced by 5-HT_2A_ receptor activation and altered K+ leak conductance; local populations represented either as explicit excitatory–inhibitory spiking networks or as mean-field models capturing their mean dynamics; and a wholebrain network linking mean-field units according to the anatomical connectivity. The bottom row shows the application of this framework to simulate both spontaneous and evoked conditions (additional external stimulus is applied to a single region) under different physiological regimes. From spontaneous activity, we extracted measures such as Lempel-Ziv complexity (LZc) and spectral features; from evoked activity, we computed the Perturbational Complexity Index (PCI).

### 2.2 5-HT_2A_ receptors and K+ conductance

Activation of 5-HT_2A_ receptors leads to closure of K+ channels (see Figure 1), resulting in neuronal depolarization, increased excitability, and a higher firing rate (see Supplementary Fig. S1). Below we show how we implement this in the model.

#### Leak Term Decomposition

The dynamics of an isopotential neuron’s membrane potential are governed by:

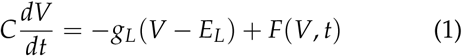

where:

- *C*: Membrane specific capacitance (F/cm^2^)
- *V*: Membrane potential (mV)
- *g*_*L*_: Leak conductance (nS)
- *E*_*L*_: Leak reversal potential (mV)
- *F*(*V, t*): Function encompassing synaptic inputs and active currents

To model the action of 5-HT_2A_ receptors, we take into account experiments showing that the stimulation of these receptors blocks leak K+ conductances in all cortical neuron types [23, 24, 25, 26, 27, 28]. To do this, we express the leak conductance *g*_*L*_ as a sum of Na+ and K+ leak conductances, *g*_*Na*_ and *g*_*K*_, such as:

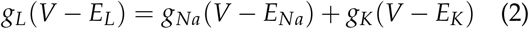

where *E*_*Na*_ and *E*_*K*_ are their respective reversal potentials.

Expanding and matching terms in (2):

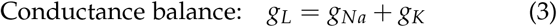

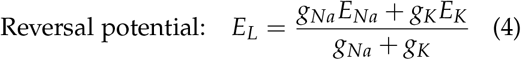

From (3) and (4), we derive:

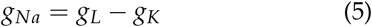

Solving (4) for *g*_*K*_:

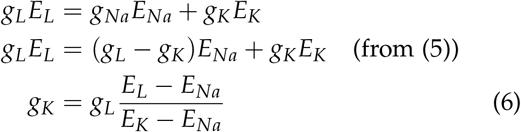

#### Baseline State

Given the following baseline values:

- *g*_*L*_ = 10 nS, *E*_*L*_ = −65 mV
- *E*_*Na*_ = 50 mV, *E*_*K*_ = −90 mV

Using (6), we compute the following:

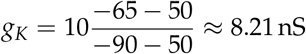

Then from (5):

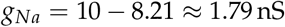

For *g*_*L*_ = 10 nS and *E*_*L*_ = −65 mV, we have *g*_*Na*_ ≈ 1.79 nS and *g*_*K*_ ≈ 8.21 nS.

#### Depolarized State

To model the effect of 5-HT_2A_ receptors, we can reduce *g*_*K*_ until the resting membrane potential reaches a desired value, here −55 mV (see Fig. 1). To compute the new *g*_*K*_ associated with this target value, we fix *g*_*Na*_ to its previously computed value (here 1.79 nS) and solve the previous system for *g*_*L*_ and *g*_*K*_:

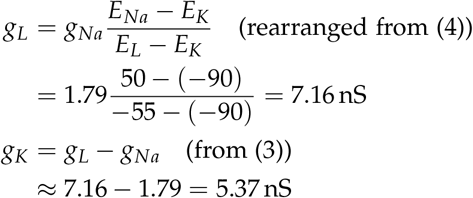

When 5-HT_2A_ receptors are activated, *g*_*K*_ decreases, leading to an increase in *E*_*L*_ and *V* and, consequently, an increase in average firing rate (see Supplementary Fig. S1).

### 2.3 Lempel-Ziv Complexity as a Marker of Consciousness in Psychoactive States

A central hypothesis in consciousness research proposes that conscious experience arises from the brain’s capacity to generate and integrate complex patterns of neural activity [48, 49]. This complexity can be quantified using measures such as Lempel-Ziv complexity (LZc), a non-parametric metric of algorithmic complexity that estimates the number of distinct subsequences in a time series, thereby indexing its information content and unpredictability [50].

In EEG and MEG studies, LZc has proven sensitive to state-dependent changes in neural dynamics, with particular relevance to serotonergic psychedelic states [51, 30, 32, 33]. Accumulating evidence indicates that these agents reliably increase LZc, reflecting heightened signal diversity and information richness during altered states of consciousness. Increases in LZc are consistently associated with subjective reports of intensified perceptual and cognitive experiences, supporting the “entropic brain” hypothesis [52, 53], which posits that the richness of conscious experience emerges from a dynamic balance between order and disorder in brain activity. Conversely, reduced-consciousness states such as deep non-REM sleep, general anesthesia, or coma are characterised by decreased LZc, consistent with synchronized, low-complexity neural dynamics [54, 55, 56, 57, 58, 59, 60, 61, 62].

While LZc effectively captures global differences in neural dynamics, interpreting its changes under psychedelics requires a mechanistic bridge from molecular-level 5-HT_2A_ receptor agonism to whole-brain effects. To address this, we developed a biophysically based whole-brain model that simulates cortical network dynamics under serotonergic modulation. We modeled the effects of reduced potassium (K+) conductance—consistent with 5-HT_2A_-mediated depolarization—on network complexity. Specifically, we set our model in two conditions, a baseline condition and a reduced K+ leak conductance described in Section 2.2 and run 15 seeds of spontaneous activity of each condition. Figure 3 (left) shows that K+ conductance reduction significantly increased simulated LZc relative to baseline, mirroring the elevated complexity reported in empirical EEG/MEG data under psychedelics. This effect was robust, as confirmed by a systematic sensitivity analysis (Figure 3, right) in which LZc differences persisted across a broad range of increased excitatory and inhibitory reversal potentials. The prevalence of significant increases across the parameter space demonstrates that the modeled LZc enhancement is not contingent on fine-tuned conditions but emerges as a general property of the network under 5-HT_2A_-like modulation. These results provide mechanistic support for the hypothesis that serotonergic psychedelics enhance cortical complexity, aligning both with empirical findings and theoretical accounts of an entropic reorganization of brain dynamics. To further probe the specificity of this effect, we examined the distribution of LZc across regions under various receptor maps (see Supplementary Figs. S2 and S3), as depicted in boxplots for baseline conditions alongside 5-HT_2A_ receptor-mediated implementations, which applied region-specific reductions in leak potassium conductance proportional to empirical 5-HT_2A_ receptor density maps.

**Figure 3:**
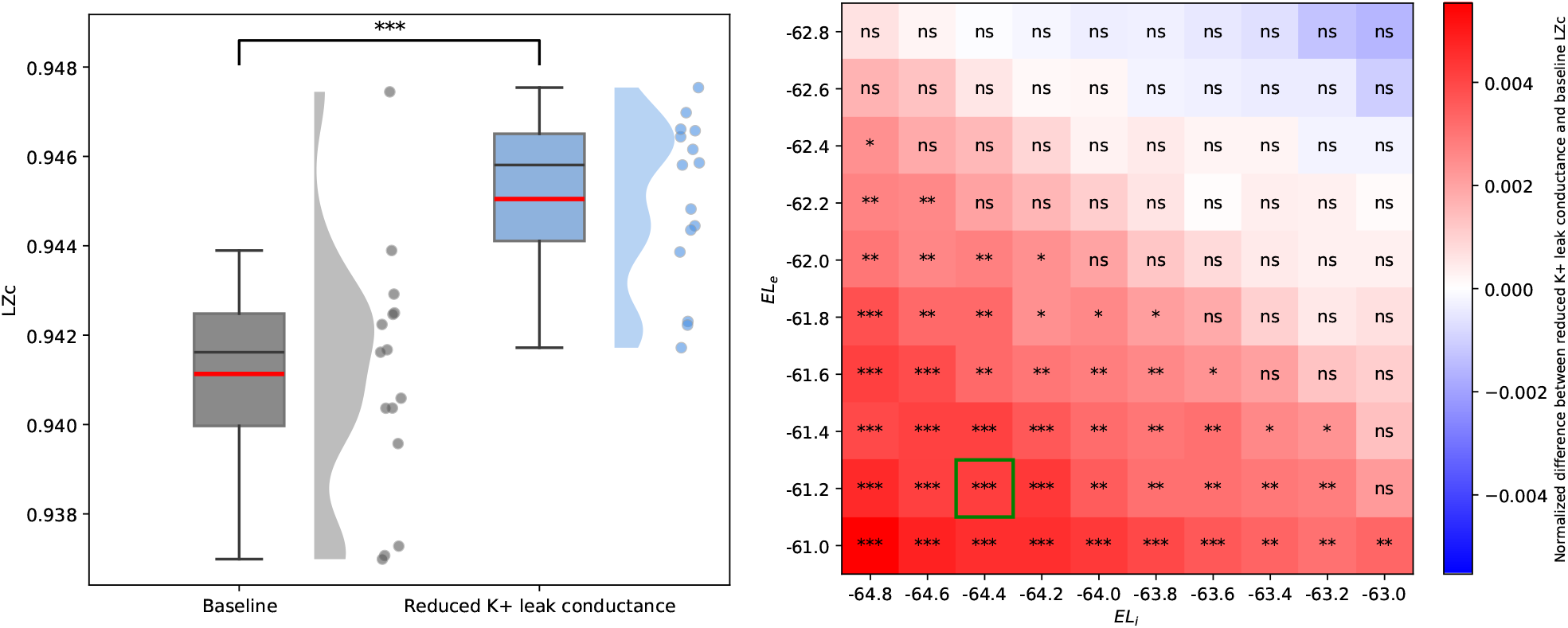
Lempel-Ziv complexity (LZc) and Robustness Analysis. Left) This figure shows the difference in Lempel-Ziv complexity (LZc) between the Baseline and reduced K+ leak conductance conditions. The box plot displays the median (red line), interquartile range (box), and whiskers extending to 1.5 times the interquartile range. The violin plot shows the data distribution, and individual data points corresponding to whole-brain simulations are overlaid. A significant difference, as indicated by the three asterisks (t-test, *p <* 0.001), is observed between the two conditions, with the reduced K+ leak conductance condition showing a higher LZc value. This is consistent with an increase in complexity reported in the literature under serotonergic psychedelics. Right) The heatmap demonstrates the robustness of the LZc results to changes in the target excitatory and inhibitory reversal potentials (baseline *EL*_*e*_ = −63 and *EL*_*i*_ = −65). The highlighted cell in green corresponds to the parameters used in the figure on the left. The color bar on the right shows the normalized LZc difference, where red indicates a positive difference (increase in LZc) and blue a negative difference. The asterisks indicate the level of statistical significance (t-tests, ∗ ∗ ∗: *p <* 0.001, ∗ ∗: *p <* 0.01, ∗: *p <* 0.05, ns: *p >* 0.05). The prevalence of significant differences across a wide range of parameter values confirms that the observed increase in LZc under the reduced K+ leak conductance condition is a robust finding.

### 2.4 Perturbational Complexity Index in Simulated Baseline and Psychedelic States

The Perturbational Complexity Index (PCI) quantifies the brain’s capacity to integrate and differentiate information in response to a controlled external perturbation [63]. It is computed from the algorithmic complexity of transcranial magnetic stimulation (TMS)-evoked electroencephalographic (EEG) responses and has been validated as a robust, theory-informed biomarker of the conscious level, independent of task demands or sensory processing. High PCI values are a hallmark of wakefulness, reflecting rich and distributed cortical interactions, whereas markedly reduced PCI values consistently characterise states of diminished consciousness, including non-rapid eye movement (NREM) sleep, general anaesthesia, and disorders of consciousness such as the vegetative state/unresponsive wakefulness syndrome (VS/UWS) [63, 64].

Recent empirical findings have revealed a notable dissociation between PCI and the subjective alterations of consciousness induced by serotonergic psychedelics: despite marked changes in perception and cognition, PCI values remain in the range typical for normal wakefulness [32]. A similar effect has been reported under low-dose ketamine, which produces a non-serotonergic psychoactive state [31]. This suggests that while psychedelics profoundly modify the content of consciousness, they do not diminish its capacity, in contrast to unconscious states. Such results highlight PCI’s specificity to global integrative capacity and motivate its combined use with spontaneous signal complexity measures—such as Lempel-Ziv complexity—to capture the full neurophysiological impact of psychedelics.

To mechanistically investigate this dissociation, we used our model in two conditions: (i) baseline wakefulness, and (ii) a psychedelic state implemented as a reduction in K+ conductance to mimic the net excitability changes induced by 5-HT_2A_ activation. In each simulation, a focal perturbation—a square wave of 0.5 Hz amplitude and 50 ms duration—was delivered to the excitatory population of the caudal right middle frontal gyrus, and PCI was computed across 15 different noise seeds. As shown in Figure 4 (left panel), the simulated psychedelic condition did not produce a statistically significant change in PCI compared to baseline, consistent with empirical TMS–EEG data in humans under psilocybin [32]. The distribution of PCI values was highly overlapping between the two conditions, indicating preserved large-scale causal integration despite the simulated neurochemical modulation. The robustness analysis (Figure 4, right panel) further confirmed that this lack of significant PCI change was stable across a wide range of excitatory and inhibitory reversal potential values, with most parameter combinations showing no significant difference (“ns”) between conditions. To ensure that the choice of stimulation amplitude did not bias PCI estimates, we systematically compared baseline and reduced *g*_*K*_ conditions across a range of stimulation amplitudes (0.25–1 Hz) and multiple seeds. These control analyses revealed no significant differences in PCI outcomes across stimulation values, thereby confirming the robustness of our results to stimulation amplitude within this range (see Supplementary Fig. S6). On this basis, we selected an amplitude of 0.5 Hz for all subsequent simulations, balancing consistency with prior work and numerical stability.

**Figure 4:**
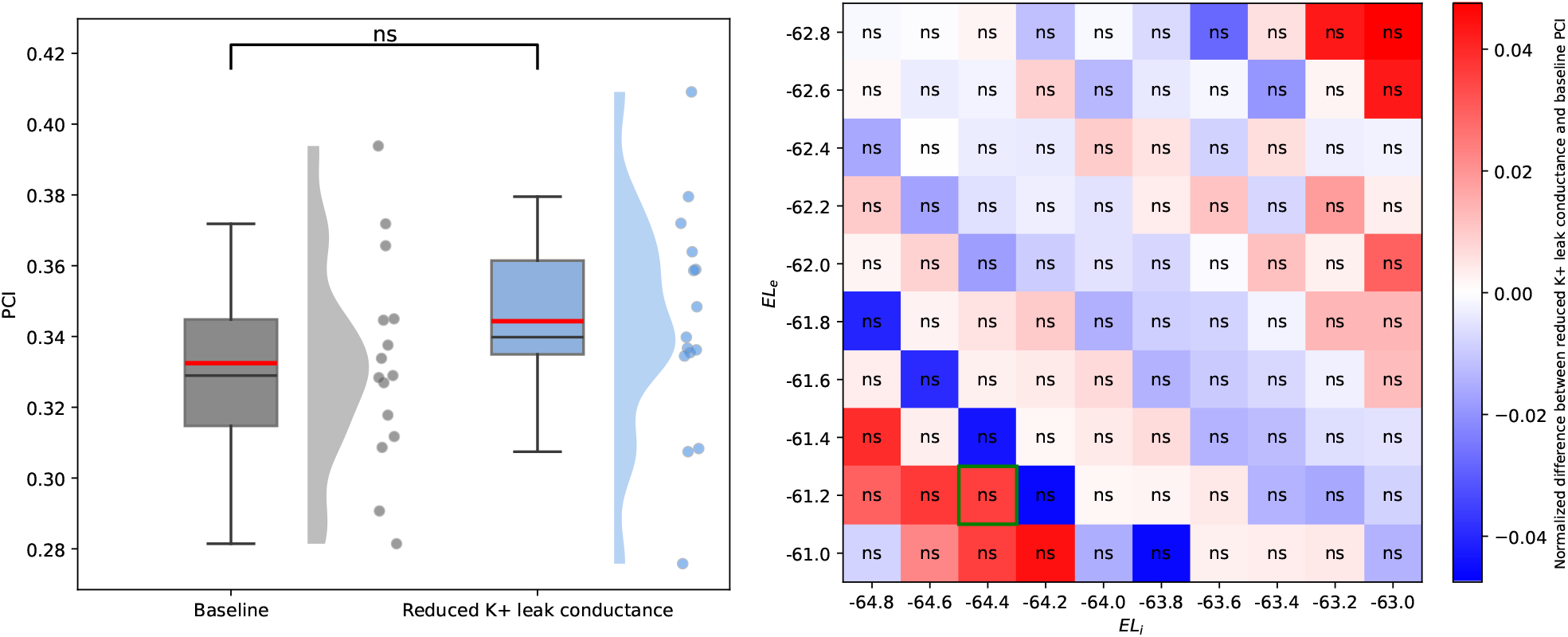
Perturbational Complexity Index (PCI) Analysis and Robustness. Left) The box plot compares the Perturbational Complexity Index (PCI) values between the Baseline and reduced K+ leak conductance conditions. The central red lines represent the median PCI values, while the boxes indicate the interquartile range (IQR). The whiskers extend to 1.5 times the IQR. The violin plots show the distribution of the simulations and individual points are overlaid. The results show no statistically significant difference (“ns”, t-test, *p >* 0.05) in PCI between the two conditions, which is consistent with findings in the literature regarding the effects of psilocybin on PCI. Right) The heatmap demonstrates the robustness of the PCI results under changes in key model parameters. The x- and y-axes represent variations in the excitatory and inhibitory target reversal potentials (baseline *EL*_*e*_ = − 63 and *EL*_*i*_ = −65). The highlighted cell in green corresponds to the parameters used in the figure on the left. The color scale represents the normalized PCI difference, with red indicating a positive difference and blue a negative difference. The abundance of “ns” (not significant) labels across the heatmap confirms that the lack of a significant change in PCI, showing the robustness of our findings across a wide range of these parameter values.

### 2.5 Spectral Power in Simulated Baseline and Psychedelic States

Alterations in spontaneous spectral power are among the most consistent electrophysiological signatures of serotonergic psychedelic states. Across EEG and MEG studies these compounds reliably reduce power in low- and mid-frequency bands—particularly in the alpha range—while often increasing spontaneous signal diversity (Table 1). This pattern has been interpreted as reflecting a breakdown of oscillatory synchrony in canonical resting-state networks, such as the default mode network, in line with theories of increased neural entropy under psychedelics [52].

**Table 1.**
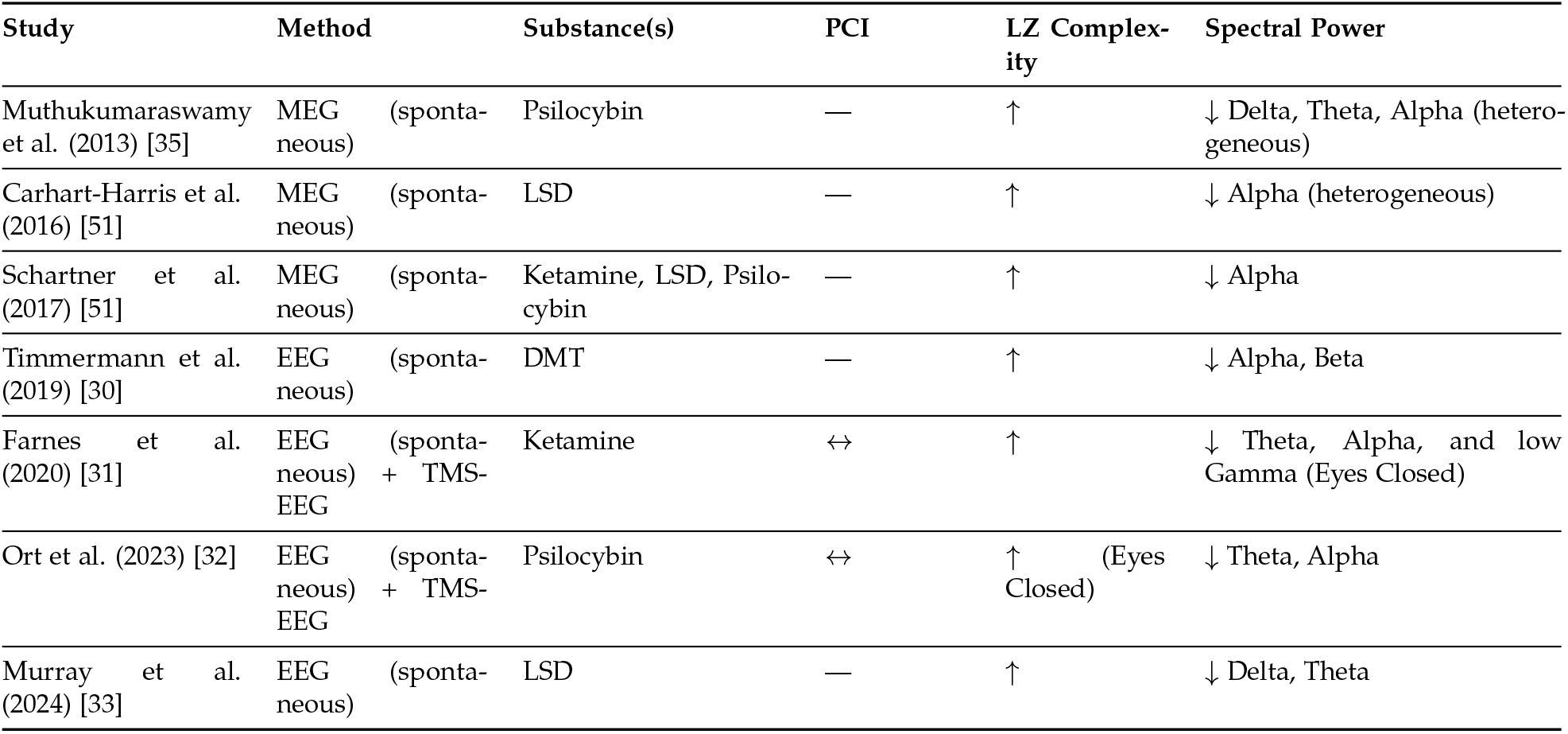
Summary of neurophysiological findings in psychedelic research. Studies are ordered chronologically. Arrows indicate: ↑ increase, ↓ decrease, ↔ no change, — not measured. LZ = Lempel-Ziv.

| Study | Method | Substance(s) | PCI | LZ Complexity | Spectral Power |
| --- | --- | --- | --- | --- | --- |
| Muthukumaraswamy et al. (2013) [35] | MEG (spontaneous) | Psilocybin | — | ↑ | ↓ Delta, Theta, Alpha (heterogeneous) |
| Carhart-Harris et al. (2016) [51] | MEG (spontaneous) | LSD | — | ↑ | ↓ Alpha (heterogeneous) |
| Schartner et al. (2017) [51] | MEG (spontaneous) | Ketamine, LSD, Psilocybin | — | ↑ | ↓ Alpha |
| Timmermann et al. (2019) [30] | EEG (spontaneous) | DMT | — | ↑ | ↓ Alpha, Beta |
| Farnes et al. (2020) [31] | EEG (spontaneous) + TMS-EEG | Ketamine | ↔ | ↑ | ↓ Theta, Alpha, and low Gamma (Eyes Closed) |
| Ort et al. (2023) [32] | EEG (spontaneous) + TMS-EEG | Psilocybin | ↔ | ↑ (Eyes Closed) | ↓ Theta, Alpha |
| Murray et al. (2024) [33] | EEG (spontaneous) | LSD | — | ↑ | ↓ Delta, Theta |

To assess whether our model reproduces these empirical spectral changes, we computed the channel-averaged power spectral density (PSD) for two simulated conditions: (i) baseline wakefulness and (ii) a psychedelic state induced by reducing K+ leak conductance to mimic the excitability effects of 5-HT_2A_ receptor activation. The PSD was estimated for each condition, averaged across 15 independent noise realizations, and decomposed into conventional EEG frequency bands (*δ, θ, α, β, γ*).

As shown in Figure 5 (left panel), the psychedelic simulation exhibited a significant (*p* < 0.001) reduction in spectral power within the *δ, θ*, and *α* bands, closely mirroring human EEG and MEG findings under psilocybin [34, 35, 32], Lysergic acid diethylamide (LSD) [51], and dimethyltryptamine (DMT) [30]. No significant changes were observed in the *β* or *γ* bands. Shaded areas represent variability across simulations, with solid lines denoting the mean PSD.

**Figure 5:**
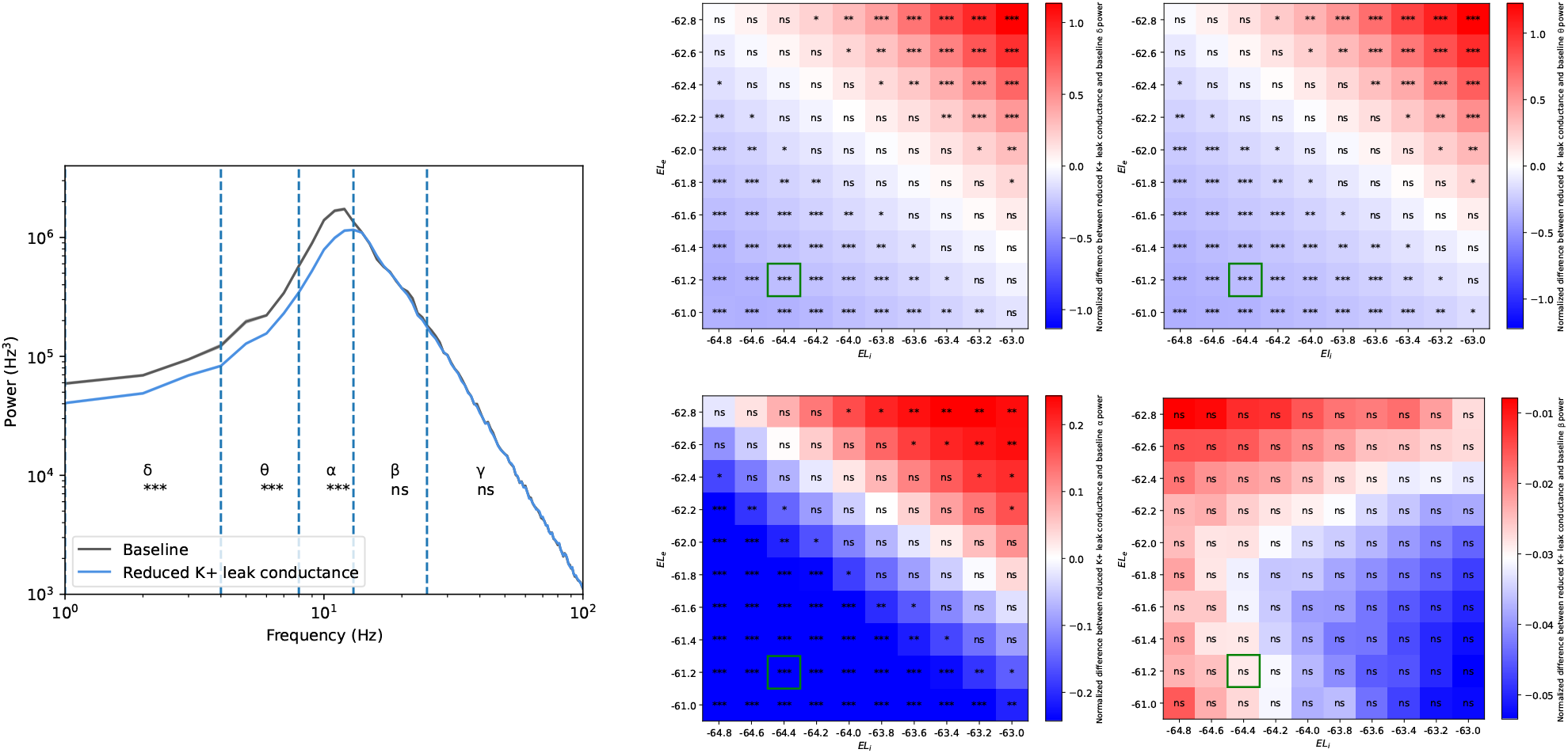
Spectral Power Analysis and Robustness to Parameter Variations. Left) The plot shows seed- and channel-average spontaneous Power Spectral Density (PSD) plotted on a semi-log scale, highlighting the difference between the Baseline (black line) and reduced K+ conductance (blue line) conditions. The PSD was computed for both conditions and segmented into conventional EEG bands for analysis (*δ, θ, α, β, γ*). The plot reveals a significant decrease in spectral power in the *δ, θ, α* bands under the reduced K+ leak conductance condition, indicated by the triple asterisks (t-test, *p* > 0.001). These results align with a known decrease in these bandwidths observed in literature under serotonergic psychedelics. Shaded areas represent the standard deviation across simulations and channels, while solid lines represent the average PSD. Right) These heatmaps illustrate the normalized differences in spectral power for the *δ, θ, α* and *β* bands, demonstrating the robustness of the observed effect. No significant difference was found in the gamma bands in any of the tested configurations. The x- and y-axes represent variations in the inhibitory and excitatory (*EL*_*i*_ and *EL*_*e*_) reversal potentials, respectively (baseline *EL*_*e*_ =− 63 and *EL*_*i*_ =− 65). The highlighted cells in green correspond to the parameters used in the figure on the left. The color gradient indicates the magnitude and direction of the normalized spectral power difference, with red signifying a positive difference and blue a negative difference (t-tests, ∗ ∗ ∗ : *p* < 0.001, ∗ ∗ : *p* < 0.01, ∗ : *p* < 0.05, ns: *p* > 0.05). This figure demonstrates that the decrease in the *δ, θ, α* channels is a robust finding that holds true across a range of reversal potential values for both excitatory and inhibitory neural populations.

The robustness analysis (Figure 5, right panel) confirmed that the power reduction in *δ, θ*, and *α* bands was not dependent on a narrow set of model parameters: across a broad range of excitatory and inhibitory reversal potential values, the effect persisted, with most parameter combinations yielding statistically significant decreases (absence of “ns” labels).

These results show that our model captures an important feature of psychedelic neurophysiology: a robust reduction in low-frequency oscillatory power, especially in the alpha band, co-occurring with increased spontaneous complexity (Section 2.3) and preserved perturbational complexity (Section 2.4). This convergence between simulated and empirical data supports the model’s validity in linking receptor-level modulation to large-scale electrophysiological changes observed in human brain recordings.

## 3 Discussion

Understanding the intricate mechanisms by which serotonergic psychedelics alter human brain activity at the whole-brain level remains a significant challenge in neuroscience. The multiscale modeling framework presented here addresses this challenge from a modeling perspective by integrating biophysical details, from single-neuron properties to large-scale network dynamics, providing a principled approach to investigate the effects of these compounds. By explicitly incorporating the action of psychedelics on 5-HT_2A_ receptors and their influence on leak membrane K+ conductances, our model offers a mechanistic bridge between molecular-level perturbations and emergent macroscopic brain states.

A key finding of this work is the model’s ability to robustly replicate the experimentally observed increase in brain complexity, as quantified by the Lempel-Ziv complexity (LZc) index, under simulated psychedelic conditions. This increase in LZc, characterised by asynchronous-irregular dynamics and enhanced complexity compared to the resting waking state, aligns with the “entropic brain hypothesis” [52]. Our results provide an important computational support for the notion that heightened neural signal diversity and information richness underpin the altered states of consciousness induced by these compounds.

Furthermore, the model accurately replicates experimental findings regarding the Perturbational Complexity Index (PCI), demonstrating that PCI increases, but not significantly under psychedelic drug administration. This crucial dissociation, where spontaneous complexity (LZc) is increased while perturbational complexity (PCI) is preserved, highlights the distinct nature of these two measures of consciousness. The model posits that psychedelics enhance local excitability without compromising the brain’s integrative causal architecture under perturbation. This suggests that while the content and experience of consciousness are profoundly modulated, the underlying capacity for consciousness, as indexed by PCI, is maintained. Such findings underscore the domain-specific sensitivity of different complexity metrics and emphasize the necessity of employing complementary measures to fully characterise altered states of consciousness, including high-order information theoretic measures [65, 66] and dynamic and emergence frameworks [67, 68, 69, 70].

Our simulations further demonstrate that the marked reduction in low-frequency spectral power (*δ, θ, α*) observed under diminished K+ leak conductance is a robust phenomenon, persisting across a wide range of excitatory and inhibitory reversal potentials. This effect resonates with empirical reports of reduced alpha and low-frequency power under serotonergic psychedelics, suggesting that alterations in intrinsic conductances may provide a mechanistic substrate for these characteristic spectral signatures.

Beyond the specific context of psychedelics, this multiscale modeling framework represents a significant advancement towards the creation of “digital twins” of the brain [71]. By allowing for the systematic investigation of how changes at the molecular and cellular levels translate into macroscopic patterns of brain activity, the framework enhances the precision and applicability of whole-brain modeling for both fundamental neuroscience and potential clinical applications. Future work will involve further validation against a broader range of experimental data, exploring the effects of different receptor sub-types and their distributions, and investigating the framework’s utility in modeling other pharmacological interventions and pathological brain states.

## 4 Current limitations and future research

While the present modeling framework provides a bottom-up bridge between receptor-level perturbations and emergent whole-brain dynamics, several limitations must be acknowledged. First, although our approach incorporates biologically realistic mechanisms such as 5-HT_2A_ receptor–mediated modulation of potassium leak conductance, the model currently focuses on a single receptor system. Psychedelic compounds act on multiple neurotransmitter receptors, including dopamine and glutamate [72, 73, 18], whose contributions remain to be explored in future extensions of this work. Incorporating these additional receptor targets, as well as their heterogeneous spatial distributions, would help to refine the generality and predictive power of the framework [74].

Second, our modeling approach relies on group-level structural connectivity data, which may not represent well individual differences in anatomy, receptor density, and pharmacokinetics. Integrating subject-specific connectomes and PET-based receptor maps could enable the development of personalized models of psychedelic action, providing new opportunities for linking emergent neurobiological features with individual variability. Such “digital twins” modeling approaches are particularly relevant for clinical translation.

Third, the current analysis emphasizes univariate measures of complexity and spectral properties. While these measures capture robust aspects of brain states induced by psychedelics, higher-order and dynamical functional interactions are likely essential to explain subjective phenomena such as ego dissolution and complex visual imagery. Incorporating recent developments in dynamic network analysis will be necessary to extend the explanatory scope of whole-brain models [75, 76, 67].

Future research may also explore non-linear dose–response relationships, as new evidence suggests that psychedelic compounds on high doses produce loss of consciousness and the emergence of brain states similar to those of unconsciousness. Extending the present framework to systematically investigate such dose-response relationships could reveal critical transitions in brain dynamics with direct relevance for dosing strategies.

## 5 Methods

We describe here the details of the multiscale whole-brain model: we start at the cellular level with the introduction of the single-cell model, then we move to the mesoscale with the construction of a neuronal network and the formulation of a mean-field model describing the mesoscopic network activity, and we end at the whole-brain scale using a large network of mean-fields interconnected using the anatomical connectome, step realized through the integration in The Virtual Brain simulator (Figure 2).

### 5.1 Single cell model

The dynamics of the single neuron are based on the adaptive exponential integrate-and-fire model (AdEx) and are described by the following equations [36]:

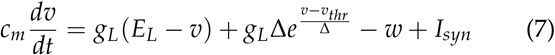

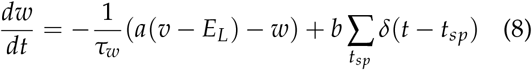

where *v* is the membrane potential, *c*_*m*_ = 200 pF is the membrane capacitance and *g*_*L*_ = 10 nS (in the baseline condition) is the leak conductance. The leakage reversal potential *E*_*L*_ equals −65 mV and -U63 mV for inhibitory and excitatory cells respectively in the baseline. The exponential term Δ has a different strength for the two types of cells (2 mV for excitatory and 0.5 mV for inhibitory cells). *w* and *I*_*syn*_ describe the adaptation and synaptic current respectively. Parameter *a* describes the sub-threshold adaptation and *b* describes the spike-triggered adaptation. A spike is generated when the membrane potential exceeds a voltage threshold *v*_*thr*_ = − 50 mV at time *t*_*sp*_. The neuron’s membrane potential is subsequently reset to a resting voltage *v*_*rest*_ = −65 mV and fixed to that value for a refractory period *T*_*refr*_ = 5 ms. The Dirac *δ*-function indicates that whenever a neuron fires at time *t*_*sp*_(*k*), the adaptation current *W* is incremented by an amount *b*. Based on physiological characteristics [77], the inhibitory cells are modeled as fast-spiking (FS) neurons, exhibiting no adaptation (*a*_*i*_ = *b*_*i*_ = 0). Excitatory neurons are modeled as regular spiking (RS) cells, characterised by lower excitability due to the presence of spiking frequency adaptation (*b*_*e*_ varies in our simulations, *a*_*e*_ = 0nS, and the adaptation time constant *τ*_*w*_ = 500ms).

### 5.2 Synaptic model

Each neuron *k* receives a synaptic current *I*_*syn*_, which corresponds to the spiking activity of all presynaptic neurons *j* ∈ pre(*k*) of neuron *k. I*_*syn*_ can be decomposed into the input received from excitatory (E) and inhibitory (I) presynaptic spikes, so as

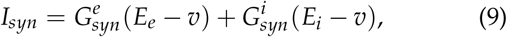

where *E*_*e*_ = 0mV (*E*_*i*_ = −80mV) is the excitatory (inhibitory) reversal potential and 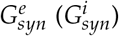 the excitatory (inhibitory) synaptic conductance. Synaptic conductances were modeled by a decaying exponential function that sharply increases by a fixed amount *Q*_*x*_, at each spiking time 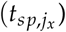 of a presynaptic neuron *j*_*x*_:

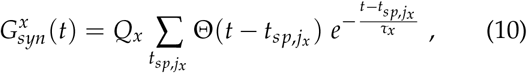

where x corresponds to the population type (*x* ∈ {*e, i*}), Θ is the Heaviside function, *τ*_*x*_ is the characteristic decay time of synaptic conductances (varied in our simulations), and *Q*_*x*_ is the quantal conductance (*Q*_*e*_ = 1.5nS, *Q*_*i*_ = 5nS). The sum runs over all the spiking times of excitatory (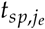) or inhibitory (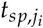) presynaptic neurons *j*_*e*_ or *j*_*i*_ of neuron *k*.

### 5.3 Spiking network model

In this work we considered a network representing a prototypical cortical circuit, specifically a single cortical column [42]. We studied a network of *N* = 10^4^ exponential-integrate-and-fire neurons that displayed spike-frequency adaptation [77, 44]. The neurons were connected over a topologically random network with a probability of connection between two neurons equal to *p* = 5%. The network was composed of two populations of inhibitory and excitatory neurons, with the inhibitory neurons consisting 20% of the whole network size.

### 5.4 Mean-field model

The mean-field used here is based on a bottom-up formalism derived from a Master Equation [43], and which has recently been extended to include the effects of adaptation [44, 38]. The mean-field equations of the system can be written as:

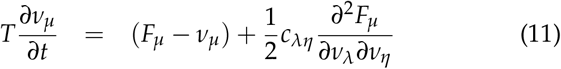

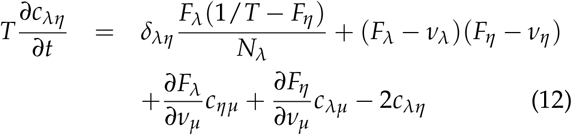

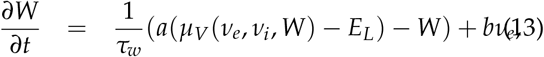

where *µ* = {*e, i*} is the neural population index (excitatory or inhibitory), *ν*_*µ*_ the mean firing rate of the corresponding population, *c*_*λη*_ the covariance between populations (*λ* and *η*), *W* is the mean adaptation, and *T* is the mean-field characteristic time constant.

The function *F*_*µ*_ = *F*_*µ*_(*ν*_*e*_, *ν*_*i*_, *W*) is the transfer function (TF) of a neuron of type *µ*, i.e. its output firing rate when receiving inhibitory and excitatory inputs with rates *ν*_*e*_ and *ν*_*i*_, and adaptation level *W*. The TF can be derived following a semi-analytic approach [44], where the output firing rate of a neuron can be written as a function of its mean subthreshold membrane voltage *µ*_*V*_, its standard deviation *σ*_*V*_, and its time correlation decay time *τ*_*V*_:

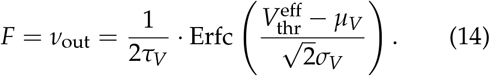

where (*µ*_*V*_, *σ*_*V*_, *τ*_*V*_) are calculated as a function of the input firing rates (*ν*_*E*_, *ν*_*I*_) and the adaptation intensity *W* following the equations described in [38]. Specifically, the mean membrane potential is calculated as the stationary solution under static conductances driven by the average synaptic input generated by firing rates (*ν*_*E*_, *ν*_*I*_). This input determines the mean 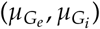 and standard deviation 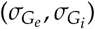 of conductances for excitatory and inhibitory processes. Assuming Poissonian spike statistics (arising from asynchronous irregular dynamics), these values are expressed as:

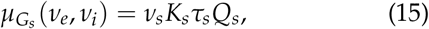

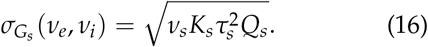

where *K*_*s*_ = *pN*_*s*_ and *s* = {*e, i*}

The mean input conductance (*µ*_*G*_) and the effective membrane time constant 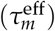 are given by:

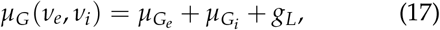

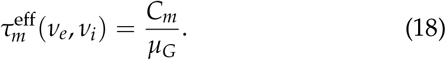

The average membrane potential for a given adaptation current *w* is:

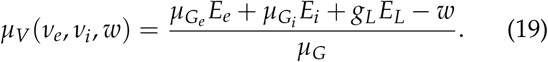

Note that in this work the terms *g*_*L*_ and *E*_*L*_ are respectively expressed by (3) and (4).

The standard deviation (*σ*_*V*_) and time constant (*τ*_*V*_) of voltage fluctuations are:

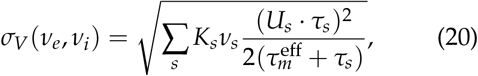

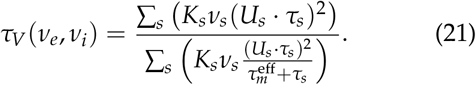

where *s* = {*e, i*} and *U*_*s*_ = *Q*_*s*_*µ*_*G*_(*E*_*s*_ *µ*_*V*_).

The 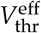 in Eq. 14 is the phenomenological spike threshold voltage taken as a second-order polynomial:

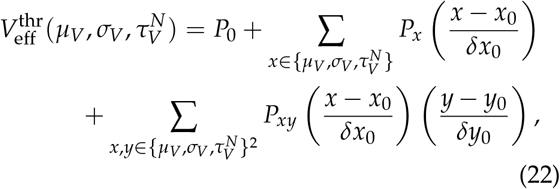

where 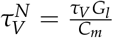 a non-dimensional quality. The polynomial coefficients of *P* are determined through a fitting of the TF template to the output firing rate of individual neuron simulations, varying both inhibitory and excitatory inputs. The values for the normalization of the fluctuation regime were set following previous work [44, 38]: 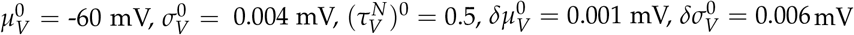 and 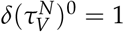. The fit is performed for each of the cell types included in the network, so in our case for fast and regular spiking cells. As shown in [38] the mean-field predictions work even far from the fitting point of the TF, so in all the examples showcased in this work we used previously calculated *P*, fitted for similar parameterizations of the two types of cells (see Table 2).

**Table 2.** (Rounded) Fitting parameters expressed in mV.

| Cell Type | $P_0$ | $P_{\mu_V}$ | $P_{\sigma_V}$ | $P_{\tau_N^V}$ | $P_{\mu_V^2}$ | $P_{\sigma_V^2}$ | $P_{\tau_N^{V^2}}$ | $P_{\mu_V\sigma_V}$ | $P_{\mu_V\tau_N^V}$ | $P_{\sigma_V\tau_N}$ |
| --- | --- | --- | --- | --- | --- | --- | --- | --- | --- | --- |
| RS-cell | -49.83 | 5.064 | -23.47 | 2.295 | -0.411 | 10.55 | -36.59 | 7.437 | 1.265 | -40.72 |
| FS-cell | -51.49 | 4.004 | -8.352 | 0.241 | -0.507 | 1.435 | -14.69 | 4.503 | 2.847 | -15.36 |

**Table 3.**
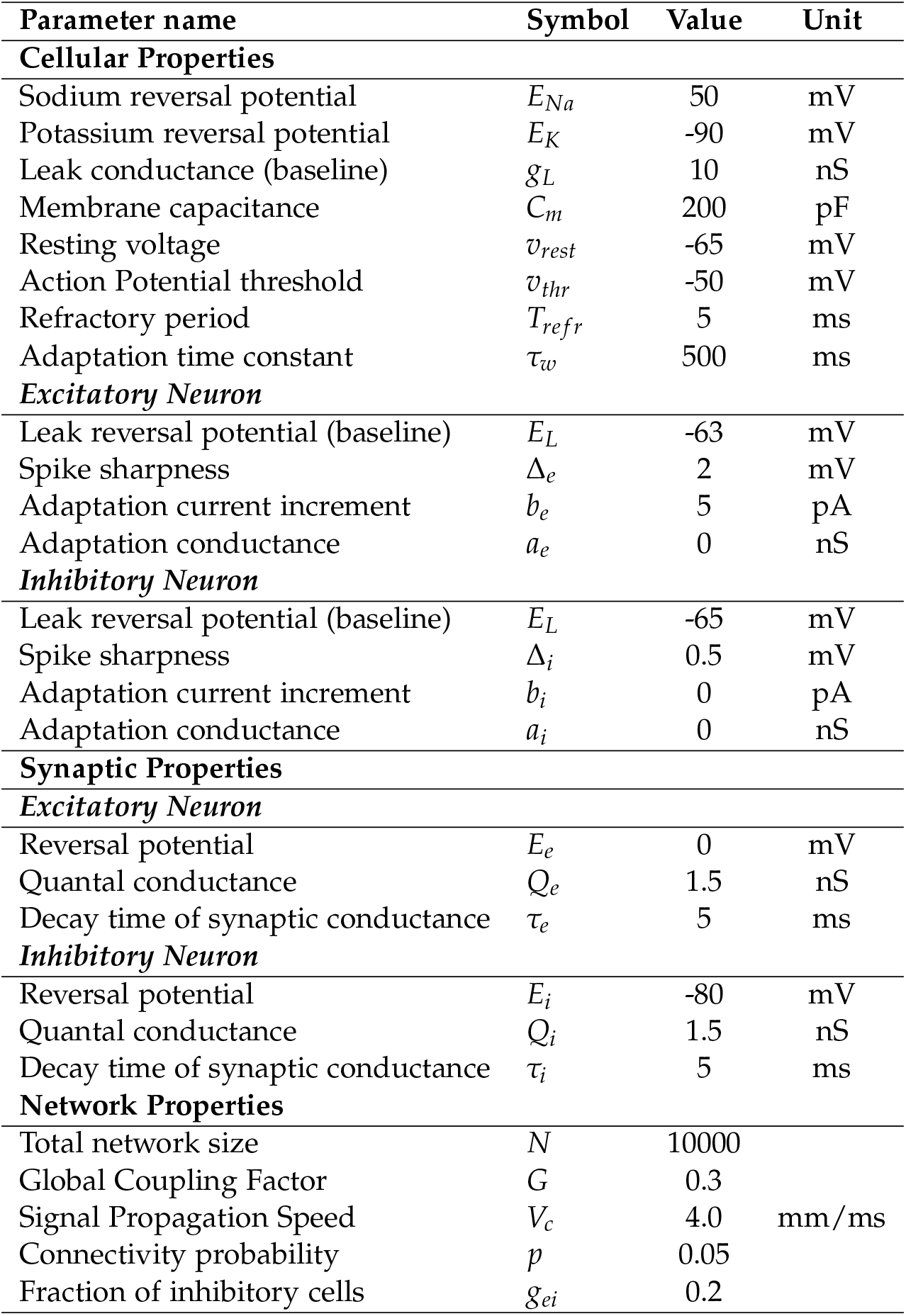
Model Parameters.

| Parameter name | Symbol | Value | Unit |
| --- | --- | --- | --- |
| **Cellular Properties** |  |  |  |
| Sodium reversal potential | $E_{Na}$ | 50 | mV |
| Potassium reversal potential | $E_K$ | -90 | mV |
| Leak conductance (baseline) | $g_L$ | 10 | nS |
| Membrane capacitance | $C_m$ | 200 | pF |
| Resting voltage | $v_{rest}$ | -65 | mV |
| Action Potential threshold | $v_{thr}$ | -50 | mV |
| Refractory period | $T_{refr}$ | 5 | ms |
| Adaptation time constant | $\tau_w$ | 500 | ms |
| **Excitatory Neuron** |  |  |  |
| Leak reversal potential (baseline) | $E_L$ | -63 | mV |
| Spike sharpness | $\Delta_e$ | 2 | mV |
| Adaptation current increment | $b_e$ | 5 | pA |
| Adaptation conductance | $a_e$ | 0 | nS |
| **Inhibitory Neuron** |  |  |  |
| Leak reversal potential (baseline) | $E_L$ | -65 | mV |
| Spike sharpness | $\Delta_i$ | 0.5 | mV |
| Adaptation current increment | $b_i$ | 0 | pA |
| Adaptation conductance | $a_i$ | 0 | nS |
| **Synaptic Properties** |  |  |  |
| **Excitatory Neuron** |  |  |  |
| Reversal potential | $E_e$ | 0 | mV |
| Quantal conductance | $Q_e$ | 1.5 | nS |
| Decay time of synaptic conductance | $\tau_e$ | 5 | ms |
| **Inhibitory Neuron** |  |  |  |
| Reversal potential | $E_i$ | -80 | mV |
| Quantal conductance | $Q_i$ | 1.5 | nS |
| Decay time of synaptic conductance | $\tau_i$ | 5 | ms |
| **Network Properties** |  |  |  |
| Total network size | $N$ | 10000 | |
| Global Coupling Factor | $G$ | 0.3 | |
| Signal Propagation Speed | $V_c$ | 4.0 | mm/ms |
| Connectivity probability | $p$ | 0.05 | |
| Fraction of inhibitory cells | $g_{ei}$ | 0.2 | |

Taking the first-order equations by disregarding the covariance term dynamics in Equation 11, the mean-field model for the excitatory and inhibitory populations representing a cortical volume can be written as:

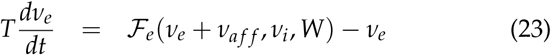

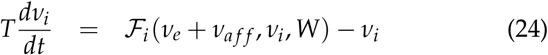

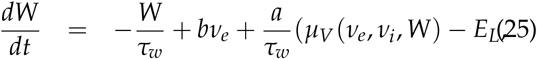

where *ν*_*a f f*_ denotes an afferent input to both type of populations which we write as:

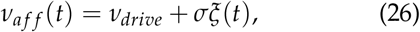

where *σ* = 3.5 and *ξ*(*t*) denotes an Ornstein-Uhlenbeck (OU) process, of the form:

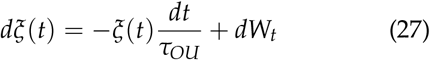

where *τ*_*OU*_ = 5ms is the timescale of the OU process and *dW*_*t*_ is a Wiener process of amplitude one and zero average. The *ν*_*drive*_ received different values in our simulations depending on the microscopic parameter that was selectively changed.

### 5.5 Networks of mean-field models

To model whole-brain dynamics, a network of mean-fields is defined, where each mean-field describes the activity of a brain region. The interactions between the mean-fields are excitatory while the inhibitory connections are preserved solely on the regional level. Extending on the single mean-field equations (Eq. 23), now the population activity of each region is given by the following equations:

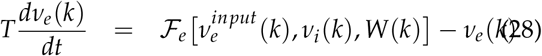

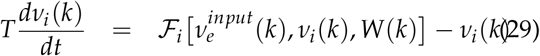

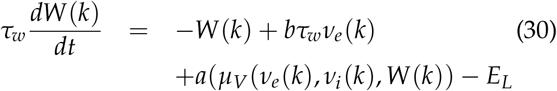

where *ν*_*e*_(*k*) (*ν*_*i*_(*k*)) describes the firing rates of the excitatory (inhibitory) population of the region (*k*), *W*(*k*) the adaptation of the population, and 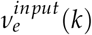 is the total excitatory synaptic input that the region receives from the rest of the nodes of the network, given by the following equation:

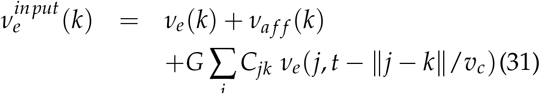

where the sum runs over all nodes *j* connected to node *k*, and *C*_*jk*_ the connection strength between *j* and *k* (and is equal to 1 for *j* = *k*), and the global coupling factor *G* rescales the connection strength while maintaining the ratio between different values. The term ∥ *j* − *k* ∥ is the distance between the nodes *j* and *k*, while *v*_*c*_ is the speed with which the signal propagates along the axis so that the model accounts for the delay of axonal propagation. Here *ν*_*a f f*_ is defined as in Eq. 26.

A cortical parcellation of 68 regions was used. The connection strengths and tract lengths between the nodes were provided by the Berlin empirical data processing pipeline, based on human tractography methods [78]. To verify robustness of the results across connectomes, the same simulations were performed using a 412 regions connectome obtained from the cortical subset of regions of a 463 regions whole-brain connectome [79, 80]. The parameters used with this connectome were the same as with the 68 regions connectome, except for the targets *EL*_*e*_ and *EL*_*i*_, as smaller differences with baseline using this connectome appeared to be equivalent to larger differences in the 68 regions connectome. Similar results were obtained and can be found in the supplementary material (see Supplementary Figs. S5, S7, S8 and S9). We performed the simulations using The Virtual Brain (TVB) platform.

### 5.6 Heterogeneity

In the whole-brain model described the only parameters differentiating one region from another are found in the connectome, that is to say the weights between each region and their relative distance. However, brain regions differ in more varied ways, for instance in receptor density, cellular composition, etc. It therefore seems necessary for whole-brain models to take into account this heterogeneity between regions. We thus use a map of 5-HT_2A_ receptor densities [74, 81] to vary the activation and the effect on *g*_*K*_ accordingly. Though this hypothesis could be refined with experimental data, we assumed a linear relationship between the density of 5-HT_2A_ receptors and the effect on the leak conductance. The leak conductance 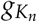 of region *n* is computed using its 5-HT_2A_ receptor density *d*_*n*_ with a linear interpolation between 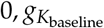 and 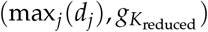, such that an hypothetical region that would have a 5-HT_2A_ receptor density of 0 would not be affected by the reduced leak conductance and the region that has the highest 5-HT_2A_ receptor density has the lowest leak conductance (and hence the highest leak reversal potential). More formally:

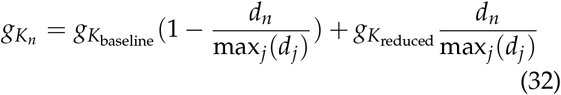

### 5.7 LZc calculation

Lempel-Ziv is a lossless compression algorithm that constructs a dictionary of substrings (words) sufficient to represent a string of characters given as input. A measure of complexity of the input string can then be obtained by simply counting the number of words that were generated in the procedure, and can be thought of as the amount of distinct patterns necessary to summarize the data. In our case, our input data are excitatory firing rates obtained from each region of the connectome in the last second of a 5 seconds simulation. This represents 10000 data points for each region. As was done in previous studies for EEG and MEG signals [82], we apply the Hilbert transform on each channel, take the absolute, and binarize the data by defining as a threshold for each channel the mean value of the channel, such that each datapoint is converted to 1 if its value is over the mean and 0 otherwise. From this binarized data, we compute two different LZc-based measures, single-channel and multidimensional. Single-channel LZc consists in averaging the channels’ individual LZc. The results of this measure on our simulations can be found in the supplementary material (Supplementary Figs. S4 and S5), and show an increase under K+ conductance reduction which holds for the two tested connectomes and multiple target leak reversal potential values, as for multidimensional LZc, which corresponds to experimental results. Multi-dimensional LZc (referred to as “LZc” in the paper) consists in first concatenating all binarized data to obtain a string of which the first *n* elements (with *n* the number of channels) are the binarized datapoints of each channel at the first timestep, the *n* following elements are the binarized datapoints of each channel at the second timestep, etc. We then apply the LZc algorithm on this binary string. For both these measures we apply a normalization scheme so that once LZc of a string is computed, we divide it by the LZc of the same string after a random permutation.

For statistical robustness we run 15 simulations with the same set of parameters but different random seeds and compute LZc each time.

### 5.8 Evoked activity and PCI calculation

The Perturbational Complexity Index (PCI), as proposed by Casali et al. [83] is the normalized Lempel-Ziv complexity of the evoked spatiotemporal patterns of cortical activation. This measure was prooposed capture the complexity of the signal propagation evoked by an external stimulation. Here, the evoked potentials were generated using the whole-brain model, where a square wave stimulus was applied on the firing rates of the excitatory population on a single node (corresponding to the right premotor cortex). 15 different trials, each with a different stimulus onset and noise realization, were performed where the analysis concerns an interval of 300 ms pre- and post-stimulus. The firing rates of the excitatory populations were first normalized using the mean and standard deviation of the pre-stimulus interval. To assess statistical significance, a bootstrap procedure was employed by repeatedly shuffling the pre-stimulus signals to generate a null distribution of surrogate values. A significance threshold was then determined from the null distribution. Finally, the normalized post-stimulus firing rates were compared to this threshold, resulting in a binary map of significant vectors *s*(*t*) for each trial. The Lempel-Ziv complexity *LZ*(*S*) was calculated for each of these vectors [9]. Finally the PCI is expressed as the ratio of 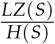 where *H*(*S*) is the spatial source entropy:

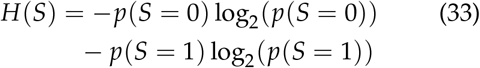

### 5.9 Power spectral density analysis

To quantify the frequency content of whole-brain activity, we computed the power spectral density (PSD) of excitatory population signals of all brain regions on the last second of 5 seconds simulations across 15 seeds for each condition. Let *x*_*r*_(*t*) denote the time series of firing rates of the excitatory population in region *r* ∈ {1, …, *N*}, sampled at frequency 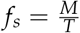, where *M* is the number of time points and *T* the total recording duration. The discrete Fourier transform (DFT) of *x*_*r*_(*t*) is given by

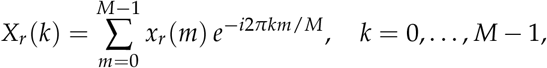

with corresponding frequencies

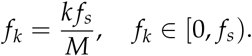

The power spectrum of region *r* is defined as the squared magnitude of the Fourier transform,

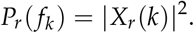

We then averaged the spectra across regions to obtain the mean PSD:

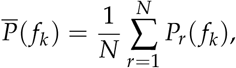

## Data availability

The connectome used in the simulations for the Desikan Killiany parcellation can be found at https://github.com/mariasacha/paper_pipeline_hub/tree/master/TVB/tvb_model_reference/data/connectivity [9]. The receptor map used in this paper for the Desikan Killiany parcellation are available at https://github.com/netneurolab/hansen_receptors [74]. The connectome and receptor maps for the Laussane 463 parcellation can be found in https://github.com/singlesp/energy_landscape/tree/v1.0.3/data [84].

## Code availability

The source code of all simulations shown in this article is available via GitHub at https://github.com/Computational-NeuroPSI/simulated_serotonergic_receptors_tvb. Our code makes use of The Virtual Brain library, available at https://www.thevirtualbrain.org/tvb/zwei (version 2.9).

## Funding acknowledgment

Research supported by the CNRS, the ANR (FLAG-ERA BrainAct project) and the European Union (Human Brain Project H2020-785907, H2020-945539, Virtual Brain Twin project 101137289, EBRAINS-2.0 project 101147319).

## Author Contributions

A.D. conceived the model, H.M. performed the analysis and simulations, R.C wrote the initial manuscript. All authors contributed to the interpretation of the results and edited the manuscript and figures. R.C and A.D supervised the project.

