## supplementary information for "Simulated 5-HT2A receptor activation accounts for the high complexity of brain activity during psychedelic states"

### Supplementary material for: Simulated serotonergic receptor stimulation reproduces the high complexity of psychedelic states at the whole-brain level

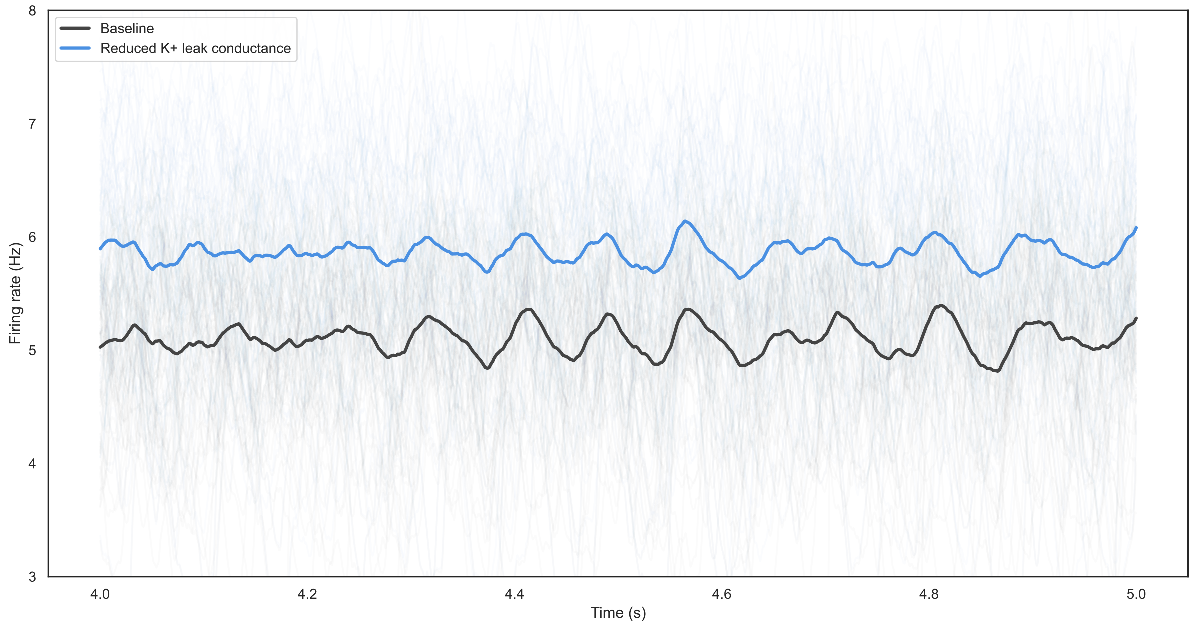


Figure S1: **Increased firing rates with reduced K+ leak conductance.** Average firing rates of excitatory populations are shown for the baseline condition (dark gray) and for the condition with reduced K+ leak conductance (light blue) where $EL_e = -61.2$ and $EL_i = -64.4$. For illustration, results are displayed for the 5th simulated second of spontaneous activity (seed 0), but the effect is robust across simulations and independent of the seed. Thin traces represent individual population firing rates, while bold lines indicate their mean dynamics over time.

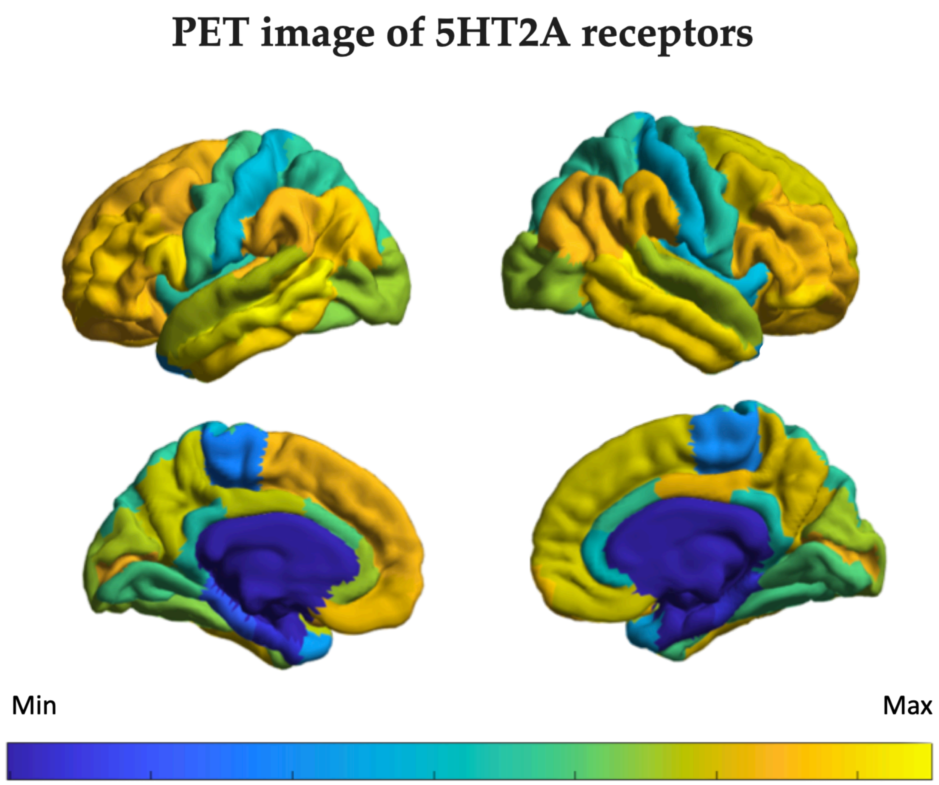


Figure S2: **5-$HT_{2A}$ Density of receptors.** The image displays a positron emission tomography (PET) derived map of serotonin 5-$HT_{2A}$ receptor density on the cortical surface using the Desikan-Killiany with 68 cortical brain areas. The color bar indicates the receptor density, with a continuous spectrum from minimum (Min) to maximum (Max) values. High receptor density is observed in regions such as the frontal cortex and posterior cingulate cortex.

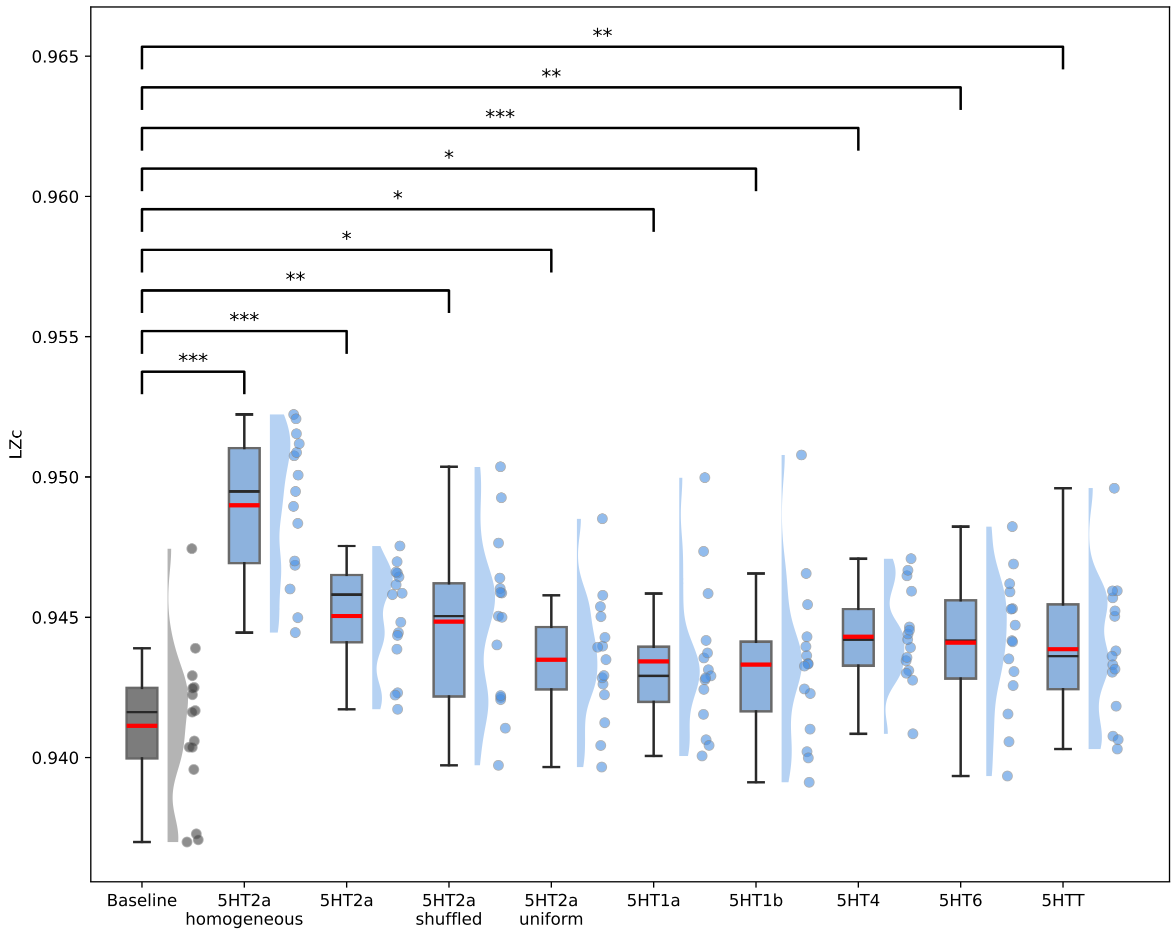


Figure S3: **Effect of receptor maps on model dynamics.** The boxplots show the distribution of the LZc across regions for different receptor maps. The baseline condition corresponds to no reduction in $g_K$. The 5-HT$_{2A}$ receptor condition implements a region-specific reduction in potassium conductance proportional to the empirical 5-HT$_{2A}$ receptor density map (as in the main simulations). Additional conditions tested the specificity of this effect: other receptor maps, a spatially randomized (shuffled) 5-HT$_{2A}$ map, uniformly distributed receptor densities sampled between the minimum and maximum values of the empirical 5-HT$_{2A}$ map (uniform), and a homogeneous map in which all regions were assigned the maximum receptor density (homogeneous). The asterisks indicate the level of statistical significance (t-tests, $***$: $p < 0.001$, $**$: $p < 0.01$, $*$: $p < 0.05$). These results show that the specific distribution of the 5-HT$_{2A}$ receptor map is not necessary to obtain a significative increase in LZc in our model.

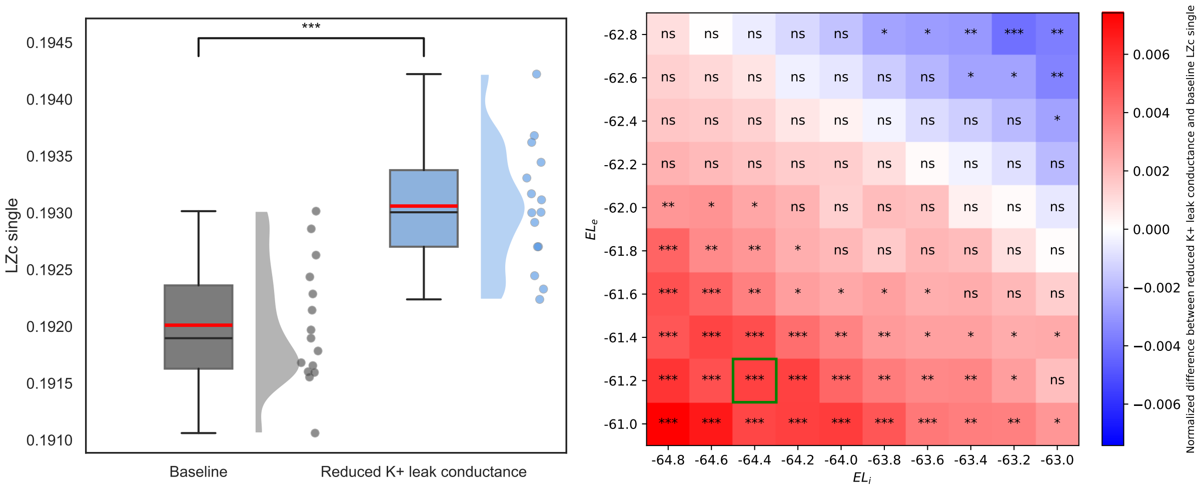


Figure S4: **Lempel-Ziv complexity for single channels (LZc single) and Robustness Analysis.** Left) Difference in LZc single between the Baseline and K+ conductance reduction conditions. The box plot displays the median (red line), interquartile range (box), and whiskers extending to 1.5 times the interquartile range. The violin plot shows the data distribution, and individual data points are overlaid. A significant difference, as indicated by the three asterisks (t-test, $p < 0.001$), is observed between the two conditions, with reduced K+ leak conductance condition showing a higher LZc single value. Right) The heatmap demonstrates the robustness of the LZc results to changes in the target excitatory and inhibitory reversal potentials (baseline $EL_e = -63$ and $EL_i = -65$). The highlighted cell in green corresponds to the parameters used in the figure on the left. The color bar on the right shows the normalized LZc single difference, where red indicates a positive difference (increase in LZc single) and blue a negative difference. The asterisks indicate the level of statistical significance (t-tests, $***$: $p < 0.001$, $**$: $p < 0.01$, $*$: $p < 0.05$, ns: $p > 0.05$). The prevalence of significant differences across a wide range of parameter values confirms that the observed increase in LZc single under the reduced K+ leak conductance condition is a robust finding.

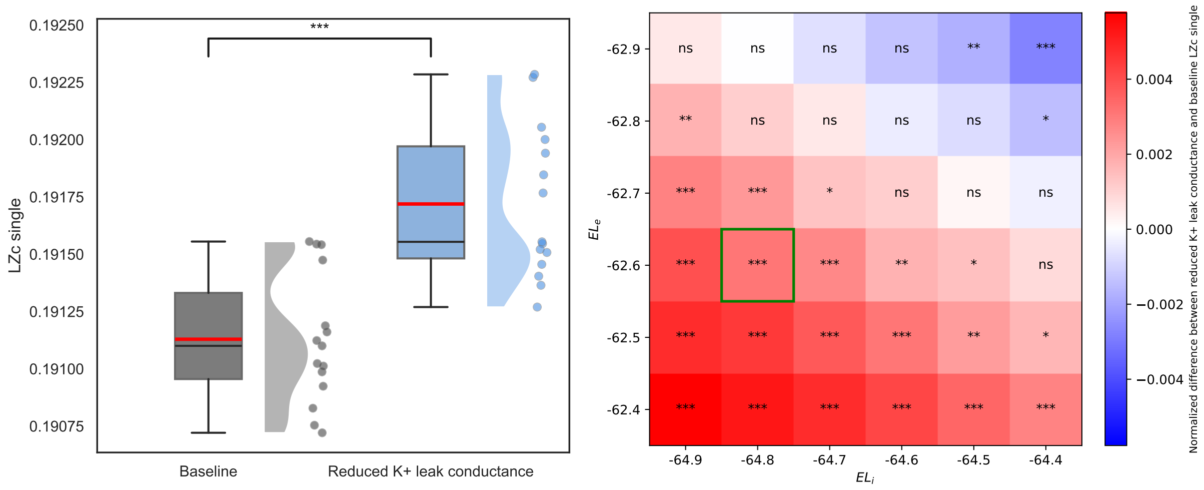


Figure S5: **Lempel-Ziv complexity for single channels (LZc single) and Robustness Analysis using the cortical-reduced Lausanne463 connectome.** Left) This figure shows the difference in LZc single between the Baseline and K+ conductance reduction conditions. The box plot displays the median (red line), interquartile range (box), and whiskers extending to 1.5 times the interquartile range. The violin plot shows the data distribution, and individual data points are overlaid. A significant difference, as indicated by the three asterisks (t-test, $p < 0.001$), is observed between the two conditions, with the K+ conductance reduction condition showing a higher LZc single value. This is consistent with an increase in complexity reported in the literature under serotoninergic psychedelics. Right) The heatmap demonstrates the robustness of the LZc results to changes in the target excitatory and inhibitory reversal potentials (baseline $EL_e = -63$ and $EL_i = -65$). The highlighted cell corresponds to the parameters used in the figure on the left. The color bar on the right shows the normalized LZc single difference, where red indicates a positive difference (increase in LZc single) and blue a negative difference. The asterisks indicate the level of statistical significance (t-tests, $***$: $p < 0.001$, $**$: $p < 0.01$, $*$: $p < 0.05$, ns: $p > 0.05$). The prevalence of significant differences across a wide range of parameter values confirms that the observed increase in LZc single under the K+ conductance reduction condition is a robust finding.

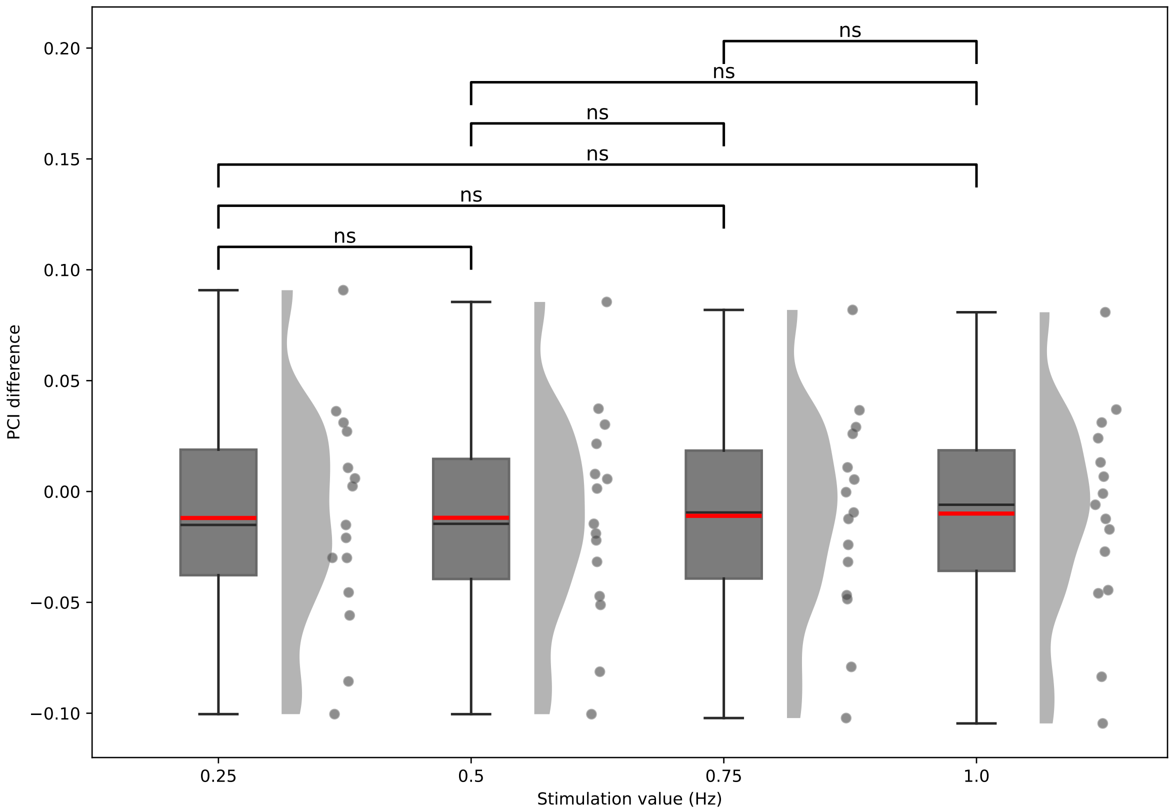


Figure S6: **Effect of stimulation amplitude on Perturbational Complexity Index (PCI)** Violin/box plots show the distribution of differences in PCI between baseline simulations (no reduction in potassium leak conductance, $g_K$) and simulations with reduced $g_K$, across multiple seeds, for a range of stimulation amplitudes (0.25–1 Hz). For each seed, the PCI difference was computed as the value under baseline minus the corresponding value under leak-reduced conditions at the given stimulation amplitude. Statistical comparisons indicate no significant differences (ns) across stimulation values, demonstrating that the choice of stimulation amplitude does not systematically influence PCI outcomes. Based on these results, a stimulation amplitude of 0.5 Hz was chosen for all subsequent analyses, ensuring comparability with previous work while avoiding instabilities (explosions) observed at higher excitatory drive (large $EL_e$ relative to $EL_i$). This control confirms that PCI findings are robust to stimulation amplitude within the tested range.

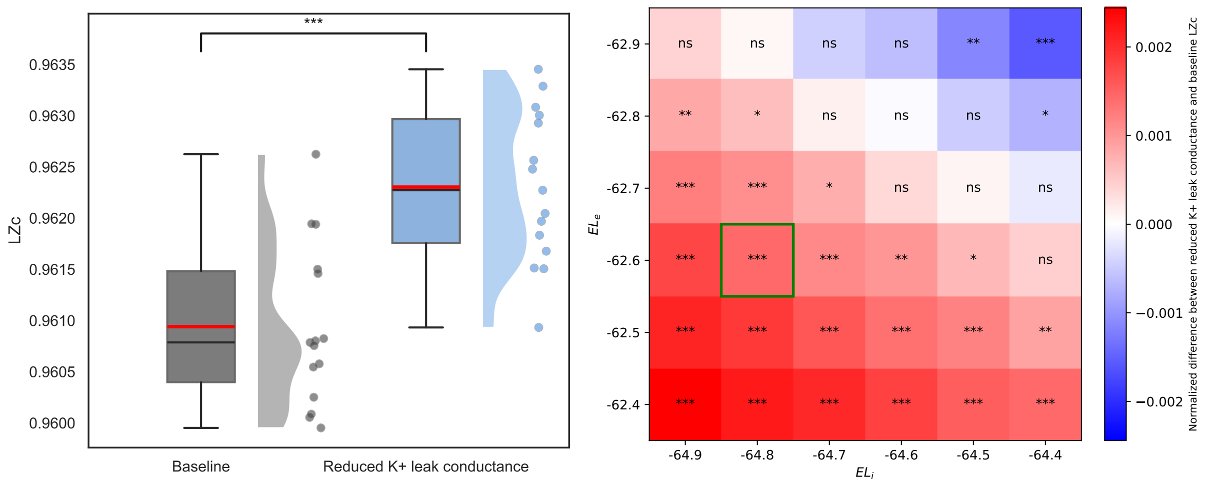


Figure S7: **Lempel-Ziv complexity (LZc) and Robustness Analysis using the cortical-reduced Lausanne463 connectome.** Left) This figure shows the difference in Lempel-Ziv complexity (LZc) between the Baseline and K+ conductance reduction conditions. The box plot displays the median (red line), interquartile range (box), and whiskers extending to 1.5 times the interquartile range. The violin plot shows the data distribution, and individual data points are overlaid. A significant difference, as indicated by the three asterisks (t-test, $p < 0.001$), is observed between the two conditions, with the K+ conductance reduction condition showing a higher LZc value. This is consistent with an increase in complexity reported in the literature under serotoninergic psychedelics. Right) The heatmap demonstrates the robustness of the LZc results to changes in the target excitatory and inhibitory reversal potentials (baseline $EL_e = -63$ and $EL_i = -65$). The highlighted cell corresponds to the parameters used in the figure on the left. The color bar on the right shows the normalized LZc difference, where red indicates a positive difference (increase in LZc) and blue a negative difference. The asterisks indicate the level of statistical significance (t-tests, $***$: $p < 0.001$, $**$: $p < 0.01$, $*$: $p < 0.05$, ns: $p > 0.05$). The prevalence of significant differences across a wide range of parameter values confirms that the observed increase in LZc under the K+ conductance reduction condition is a robust finding.

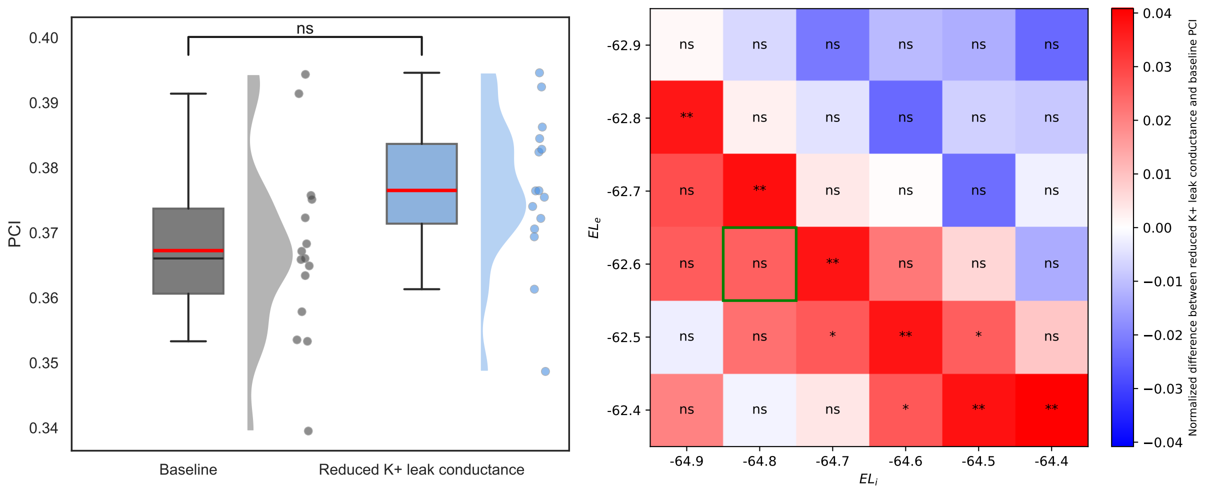


Figure S8: **Perturbational Complexity Index (PCI) Analysis and Robustness using the cortical-reduced Lausanne463 connectome.** Left) The box plot compares the Perturbational Complexity Index (PCI) values between the Baseline and K+ conductance reduction conditions. The central red lines represent the median PCI values, while the boxes indicate the interquartile range (IQR). The whiskers extend to 1.5 times the IQR. The violin plots show the distribution of the simulations and individual points are overlaid. The results show no statistically significant difference (”ns”, t-test, $p > 0.05$) in PCI between the two conditions. Right) The heatmap demonstrates the robustness of the PCI results under changes in key model parameters. The x- and y-axes represent variations in the target excitatory and inhibitory reversal potentials. The highlighted cell corresponds to the parameters used in the figure on the left. The color scale represents the normalized PCI difference, with red indicating a positive difference and blue a negative difference. The abundance of ”ns” (not significant) labels across the heatmap confirms that the lack of a significant change in PCI is a robust finding that remain true across a wide range of these parameter values.

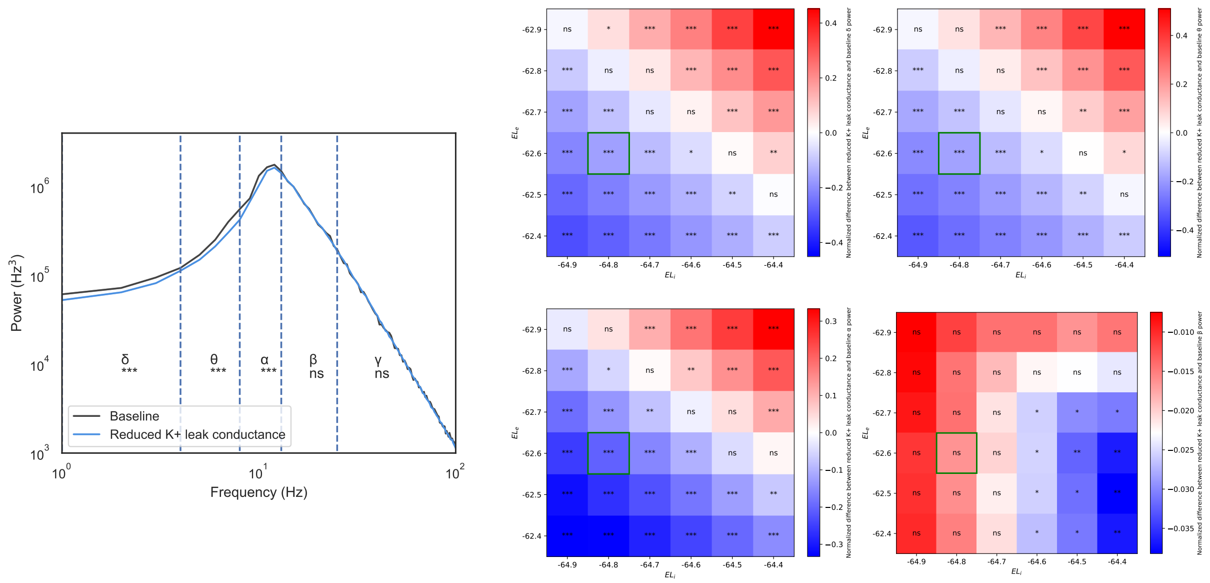

Figure S9: **Spectral Power Analysis and Robustness to Parameter Variations using the cortical-reduced Lausanne463 connectome.** Left) The plot show channel-average spontaneous Power Spectral Density (PSD) plotted on a semi-log scale, highlighting the difference between the Baseline (black line) and K+ conductance reduction (blue line) conditions. The PSD was computed for both conditions and segmented into conventional EEG bands for analysis ($\delta, \theta, \alpha, \beta, \gamma$). The plot reveals a significant decrease in spectral power in the $\delta, \theta, \alpha$ bands under the K+ conductance reduction condition, indicated by the triple asterisks (t-test, $p > 0.001$). Shaded areas represent the standard deviation across simulations and channels, while solid lines represent the average PSD. Right) These heatmaps illustrate the normalized differences in spectral power for the $\delta$, $\theta$, $\alpha$ and $\beta$ bands, demonstrating the robustness of the observed effect in the cortical-reduced Lausanne463 connectome. No significant difference was found in the gamma bands in any of the tested configurations. The x- and y-axes represent variations in the inhibitory and excitatory ($El_i$ and $El_e$) reversal potentials, respectively. The highlighted cells correspond to the parameters used in the figure on the left. The color gradient indicates the magnitude and direction of the normalized spectral power difference, with red signifying a positive difference and blue a negative difference (Student's t-tests, $***$: $p < 0.001$, $**$: $p < 0.01$, $*$: $p < 0.05$, ns: $p > 0.05$). This figure demonstrates that the decrease in the $\delta$, $\theta$, $\alpha$ channels is a robust finding that holds true across a range of reversal potential values for both excitatory and inhibitory neural populations using the cortical-reduced Lausanne463 connectome.
